# Reconciling Qualitative, Abstract, and Scalable Modeling of Biological Networks

**DOI:** 10.1101/2020.03.22.998377

**Authors:** Loïc Paulevé, Juraj Kolčák, Thomas Chatain, Stefan Haar

## Abstract

Predicting the behaviors of complex biological systems, underpinning processes such as cellular differentiation, requires taking into account many molecular and genetic elements for which limited information is available past a global knowledge of their pairwise interactions. Logical modeling, notably with Boolean Networks (BNs), is a well-established approach which enables reasoning on the qualitative dynamics of networks accounting for many species. Several dynamical approaches have been proposed to interpret the logic of the regulations encoded by the BNs. The synchronous and (fully) asynchronous ones are the most prominent, where the value of either all or only one component can change at each step. Here we prove that, besides being costly to analyze, these usual interpretations are not adequate to represent quantitative systems, being able to both predict spurious behaviors and miss others. We introduce a new paradigm, the Most Permissive Boolean Networks (MPBNs), which offer the formal guarantee not to miss any behavior achievable by a quantitative model following the same logic. Moreover, MPBNs significantly reduce the complexity of dynamical analysis, enabling to model genome-scale networks.

---

Models in systems biology typically integrate knowledge and hypotheses on molecular interactions, manually or semi-automatically, gathered from experimental data found in databases and the literature. These models are often qualified as “mechanistic,” in opposition to those solely based on biophysical laws.

Since their introduction in the late ‘60s (1, 2), logical models, such as Boolean Networks, have been widely adopted for reasoning about signaling and gene networks (3–11) as they require few parameters and can easily integrate information from omics datasets and genetic screens. These models represent processes with a high degree of generalization and can offer coarse-grained but robust predictions. That makes them particularly suitable for large biological networks, for which ample global knowledge exists about potential interactions with little precise data on actual molecules abundances and reaction kinetics.

The validation of computational models is necessary to trust their subsequent predictions. In systems biology, validation primarily involves *in silico* reproduction of observed behaviors by executing the computational model. Such observations may be measurements of the activity, over time, or at steady-state, of some of the interacting molecules under different experimental conditions. Therefore, if no executions of a BN reproduce an experimentally observed behavior (e.g., the activation of a particular gene), the model, and the associated interactions, is considered as invalid. This procedure also enables general studies on interaction motifs that are necessary or sufficient for achieving fundamental behaviors such as cellular differentiation or homeostasis (12–15).

Boolean Networks are often created from scratch and rather than derived from a detailed mechanistic (partially-parameterized) model. Consequently, there is no guarantee that their analysis can be relevant for a more precise model, and thus for the actual biological system.

Fig. 1 illustrates this issue with the incoherent feed-forward loop of type 3, I3-FFL(16). An input node 1 directly inhibits the output 3, but indirectly activates it via node 2. Theoretical studies (**? ?**) and experimental data from synthetically designed circuits (**?**) show that a monotonic activation of the input can lead to a transient activity of the output. However, it is impossible to reproduce this behavior with usual interpretations of BNs, including synchronous and asynchronous: if 1 is not active, neither 2 nor 3 can be activated. If 1 is active, 2 is active, but any transient activation of 3 is prevented (Fig. 1(d)).

**Fig. 1.**
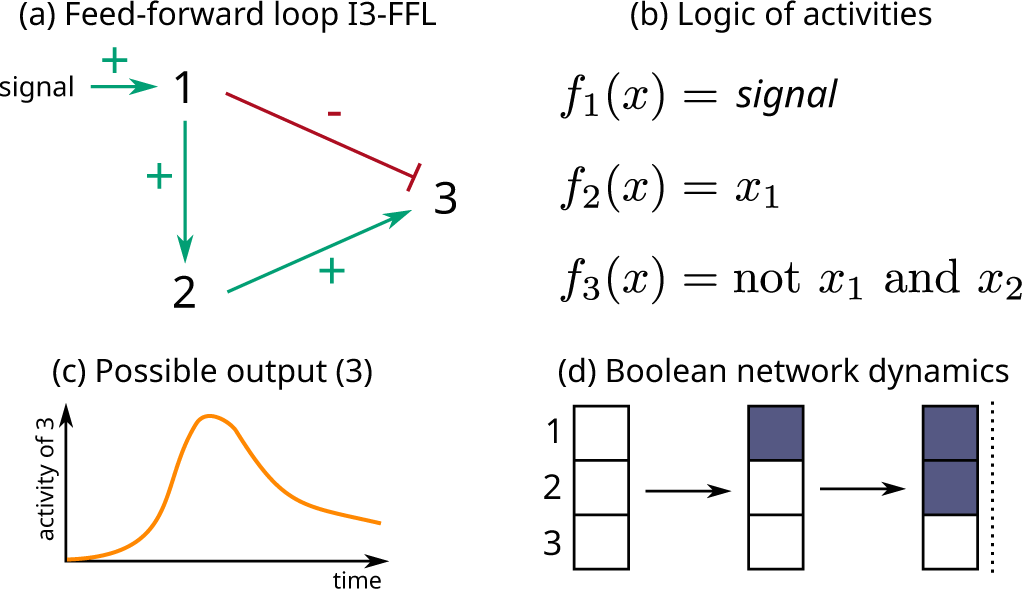
Incoherent feed-forward loop of type 3 (a) and its associated Boolean logic for nodes activities (b); *f*_1,2,3_(*x*) are the Boolean functions used to compute the next value of each node from a given *configuration x* of the network, which is here a binary vector specifying the current value of each node, *x*_*i*_ referring to the Boolean value of node *i*. Whereas theoretical and experimental studies show a possible activation of the output when the signal is active (c), usual BNs analysis cannot predict this transient behavior: (d) shows the corresponding complete dynamics of *f*, where configurations are represented by piles of 3 squares, where the top square represents the state of the first component, and so forth. A white square represents the inactive (0) state; a blue square represents the active (1) state; a dashed line indicates that no further evolution is possible. Arrows indicate possible transitions. The node 3 is never predicted to be active.

Additional model features, such as intermediate levels for the nodes, or delays in interactions, would allow a transient activation for the I3-FFL output. However, such features come with additional parameters and higher computational cost, which limits their general application to large scale networks.

This simple example seems to show that setting binary activities for nodes can both generate spurious behaviors (as expected with qualitative models), and also preclude the discovery of existing behaviors. Therefore, the validity of a model cannot be assessed by the usual interpretations of BNs. This limitation largely impedes the inference of dynamical network models and the identification of necessary interaction network motifs since the Boolean interpretation can strongly distort the landscape of candidate models.

However, we found that this issue is actually due to the usual interpretations of BNs and not to their intrinsic Boolean nature. We introduce a new simulation approach, the *Most Permissive Boolean Networks* (MPBNs), which presents the formal guarantee to capture all behaviors achievable without the need for additional parameters. If MPBNs cannot reproduce a given observation, no quantitative refinement of the Boolean model can do it, and the model can safely be considered as incoherent with the observations. While predicting more behaviors than the usual interpretations of Boolean Networks, MPBNs still capture essential dynamical features of biological models.

Moreover, we demonstrated that the analysis of MPBNs avoids the state space explosion problem, a strong limiting factor for the usual interpretations of BNs. The drastically reduced computational cost enables the precise qualitative analysis of dynamics of genome-scale networks.

## Modeling with Boolean Networks

Computational modeling of dynamical systems relies on two fundamental ingredients: a language to specify the model, and an execution semantics. The language provides symbols and syntax rules to write a model, while the execution semantics mathematically defines how to interpret it. The semantics formalizes the notion of network configurations (or states) and how to compute their evolution over time. It provides an exhaustive assessment of model capabilities by enabling dynamical analyzes such as simulations as well as formal verification by invariant analysis and model-checking.

A BN is specified by a mathematical function mapping any binary vector of dimension *n* to another binary vector of the same dimension:

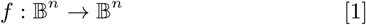

where 𝔹 = {0, 1} represent the Boolean values. Each element of a binary vector models the state (inactive/ active, absent/present) of the associated network node, and *f*_*i*_ is the function which specifies the state towards which the *i*-th element evolves. Fig. 2(b) gives an example of a BN modeling a switch system.

**Fig. 2.**
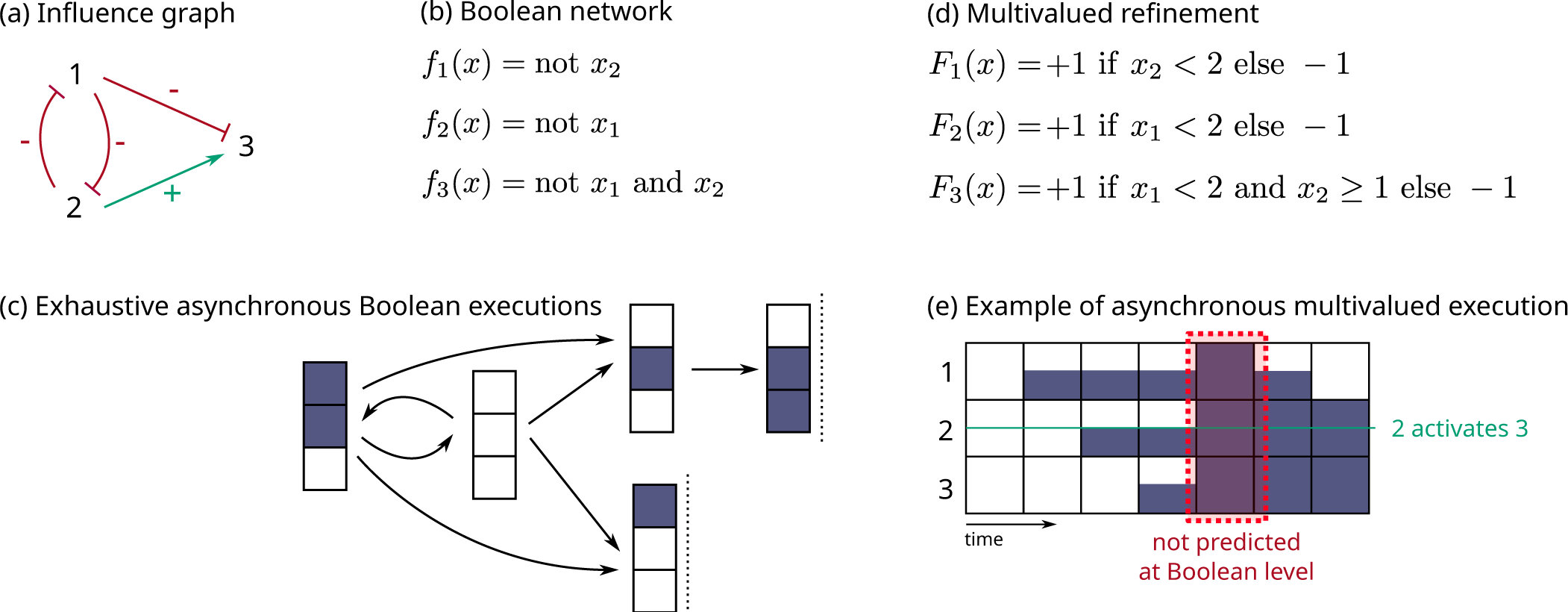
Example of qualitative models for the interactions between three components. (a) Influence graph denoting the activation and inhibition relationships. (b) Example of a compatible BN, which defines the activation conditions of each component. (c) Exhaustive list of transitions obtained from the initial configuration where all three components are inactive. (d) Example of a multivalued network refining the BN (b) with components able to exist under three states (0, 1, 2). (e) Example of asynchronous execution of the multivalued network from the configuration 000. Half and fully blue squares represent the states 1 and 2.

BNs semantics computes the possible temporal evolutions of the component states using different methods. With *synchronous* executions of BNs (introduced by S. Kauffman (1)), we update all the components of the network at the same time, and a configuration *x* ∈ 𝔹^*n*^ can only evolve to one configuration *f* (*x*). With *fully asynchronous* executions of BNs (introduced by R. Thomas and usually referred to more simply as *asynchronous* in the computational systems biology literature), we update only one component at a given time, and a configuration *x* ∈ 𝔹^*n*^ can evolve to any configuration which differs only by a single component *i* where *f*_*i*_(*x*) ≠ *x*_*i*_. This introduces potential *non-determinism* in the model trajectory since there can be different executions of the same BN from a given initial configuration. The (fully) asynchronous semantics is often described as more realistic for modeling biological networks, accounting for different kinetics of interactions.

Many more variants of executions of BNs have been studied in the literature, some imposing a precise order in the updating of the components, others allowing subsets of components to be updated simultaneously, etc. Most, if not all, generate a subset of the executions achievable with the (generalized) *asynchronous* semantics of BNs where any number of components can be updated at a time: a configuration can evolve to any other configuration that complies with the logical functions for the components that differ between both. Formally, for any *x, y* ∈ 𝔹^*n*^,

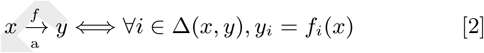

where Δ(*x, y*) is the list of components which state differs between *x* and *y*, i.e., Δ(*x, y*) = {*i* ∈ {1, …, *n*} | *x*_*i*_ ≠ *y*_*i*_}.

A configuration *y* ∈ 𝔹^*n*^ is *reachable* from *x* ∈ 𝔹^*n*^ if either *x* = *y*, or there exists a sequence of transitions from *x* to *y*:

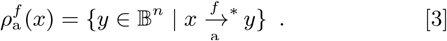

Notice that if 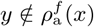, then it is impossible to evolve from *x* to *y* according to any of the semantics defined above, including the synchronous and (fully) asynchronous ones.

Fig. 2(c) shows all possible asynchronous evolutions of the example BN from the configuration where all the components are inactive, i.e. 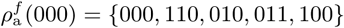.

Reachability is a fundamental property to assess the compatibility of BN models with time series data: if none of the configurations matching an observation at a given time is reachable from any configuration matching an experimental observation at an earlier time, the BN cannot predict the observed behaviors.

Another prominent dynamical property studied with BNs, strongly linked to reachability, are attractors. Attractors represent the long-term behaviors of the model and are often used to represent cell phenotypes. Formally, an attractor is a smallest non-empty set of configurations from which it is impossible to escape: *A* ⊆ 𝔹^*n*^ is an attractor if and only each of its configuration *z* ∈ *A* verifies 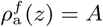. An attractor is said to be a fixed point whenever it is a single configuration *z* ∈ 𝔹^*n*^ (whenever *f* (*z*) = *z* with the asynchronous semantics), and *complex* if it is an ensemble of configurations, such as *cyclic* attractors, modeling potential sustained oscillations.

## Refinements of Boolean Networks

Boolean Networks impose a drastic coarse-graining on component activity. Several modeling frameworks introduced a finer granularity in logical models (17). Examples include Multivalued Networks (MN) (18), where components can take more than two *logical* values (0, 1, 2, …, *m*), fuzzy logic (19), which extends logical models with continuous domains, stochastic extensions of fully asynchronous Boolean Networks (20), and Ordinary Differential Equations (ODEs) (21, 22), where values of components are non-negative reals and vary along continuous time. Their specifications require, however, much more information about the biological system, such as thresholds of interactions for MNs and precise kinetics for ODEs. These parameters are often unknown, and their automatic inference would require a significant amount of data collected in similar experimental settings.

One could use any of these frameworks to model the same biological system at different abstraction levels. Which raises the question of the relationship between models from different frameworks: is an MN model *F* a refinement of a BN model *f*? In other words, does *F* specify a system with more quantitative information than *f*, but follows the same (Boolean) logic for the interactions.

We consider here a simple mathematical criterion for refinements: the value of a component can decrease (resp. increase) only if the component can be set to 0 (resp. 1) in the BN with a possible binarization of the state. A formal definition will be given in the next section.

### Incompleteness of (a)synchronous Boolean networks

We can define a multivalued network *F* of dimension *n* by a discrete function, which maps, for each component, states to the tendency of value change (decrease, steady, increase). To ease notations, and without loss of generality, we assume that all the components can take an integer value between 0 and the same fixed *m*:

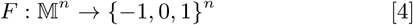

where 𝕄 = {0, 1, …, *m*}. The successors of a configuration *x* ∈ 𝕄^*n*^ are then computed by adding the value of *F* (*x*) to (a subset of) components, provided they stay non-negative and do not exceed their maximum value *m*.

In Fig. 2 we present a simple example of BN for which asynchronous executions miss possible behaviors of the network when considering a multivalued refinement of it. The MN in Fig. 2(d) is a refinement of the BN in Fig. 2(b). In addition to the higher granularity for the activity levels of all three components, it brings additional information on the activation of component 3. An intermediate value of 2 is sufficient to activate 3 provided that the value of its inhibitor 1 is not high. One of its asynchronous execution shown in Fig. 2(e) predicts that the three components can get activated simultaneously, which was never predicted by any of the asynchronous executions of the BN. Assuming the validation of the model were subject to the reachability of a configuration with all the three components active from a configuration with all the components inactive, this BN model would be deemed insufficient for achieving the observed behavior, with an erroneous conclusion that its underlying influence graph is wrong.

## A new execution paradigm for Boolean networks

The critical reason usual BN interpretations miss behaviors is that the binary coarse-graining coupled with the instantaneous state changes preempt interactions occurring *during* the course of (de)activations. In the counterexample of Fig. 2, Boolean interpretations exclude the activation of component 3 during the activation of components 1 and 2, whereas, in a possible refinement, 3 can indeed increase before 2 reaches its fully active state and before 1 is sufficiently expressed to inhibit it.

### The Most Permissive semantics

We devised a new dynamical interpretation of BNs, called Most Permissive semantics, in which we consider that a component can exist in 4 states: inactive (0), increasing (↗), decreasing (↘), or active (1). While a component is in an dynamic state (increasing or decreasing), it can be read non-deterministically as either 0 or 1. These ambiguous states account for the absence of information on actual influence thresholds: a component in a dynamic state can be above the influence threshold for one component while being below the influence threshold for another one.

Fig. 3 illustrates the changes of component states possible with the Most Permissive semantics. A component *i* can change to the increasing (resp. decreasing) state whenever it can interpret the value of its regulators in a way which makes its logical function *f*_*i*_ true (resp. false) – if one of its regulators is in a dynamic state, both Boolean interpretations can be considered. Once in increasing (resp. decreasing) state, it can reach 1 (resp. 0) at any time. Each component evolves independently of all others. The complete formal definition is given in SI.2.

**Fig. 3.**
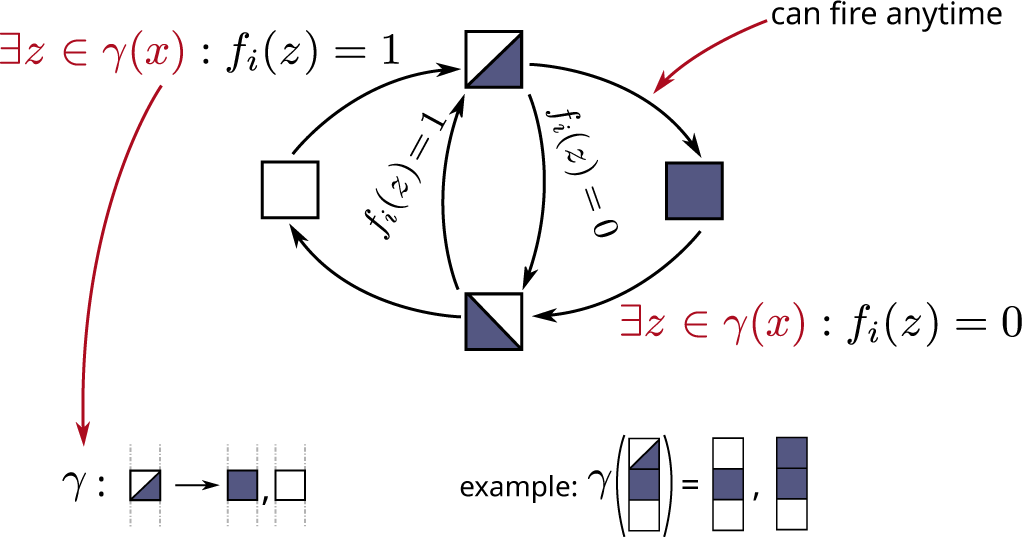
Conditions for the changes of component states *i* ∈ {1, …, *n*} in Most Permissive Boolean Networks from a most-permissive configuration *x* ∈ 𝕡^*n*^ with 𝕡 = {0, ↗, ↘, 1}. The increasing state ↗ is represented by a top-left white and bottom-right blue square, the decreasing state ↘ by a bottom-left blue and top-right white square. The function *γ* gives the admissible Boolean interpretations of *x* ∈ 𝕡^*n*^: *γ*(*x*) = {*z* ∈ 𝔹^*n*^ | ∀*i* ∈ {1, …, *n*}, *x*_*i*_ ∈ 𝔹 ⇒ *z*_*i*_ = *x*_*i*_}, *i*.*e*., all the components in Boolean states are fixed, and the others are free.

Fig. 4 shows an example of execution using the Most Permissive semantics on the BN of Fig. 2. Contrary to the (a)synchronous interpretations, the Most Permissive semantics correctly captures the possible (transient) reachability of the configuration where the three genes are active. While component 1 is “increasing” and component 2 is active, gene 3 can indeed change to “increasing”, thus leading to the activation of all three components. This configuration is not in an attractor, and both single-point attractors identified in Fig. 2(c) are reachable via different Most Permissive executions. We provide the application of MPBNs to the BN presented in Fig. 1 in in Fig. S1.

**Fig. 4.**
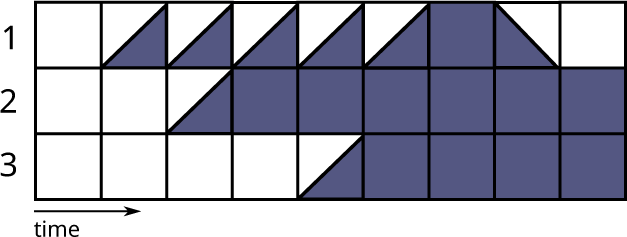
One of the possible executions using the Most Permissive semantics on the Boolean network in Fig. 2(b) starting from the configuration where all genes are inactive. Note that it correctly recovers the (transient) reachability of the configuration where the three components are active.

### Formal guarantees for model refinements

Using the simple examples in Fig. 1 and Fig. 2, we have shown that BN refinements can introduce behaviors that cannot be captured with classical semantics.

Most Permissive Boolean Networks bring the formal guarantee of being able to reproduce *all* the behaviors achievable in *any* refinements, being a multivalued network or an ODE system (Theorem 1 and Corollary 1 in SI.2). In other words, if the Most Permissive semantics concludes that it is impossible to observe a given state change for some components, then no qualitative or quantitative model verifying the refinement criteria can predict these state changes.

The refinement criterion relies on a *binarization* of the multivalued configuration. An appropriate binarization necessarily quantifies 0 as Boolean 0 and *m* as 1, and is free for the other intermediate values. Let us denote by *β* (*x*) the set of possible binarization of configuration *x* ∈ 𝕄^*n*^:

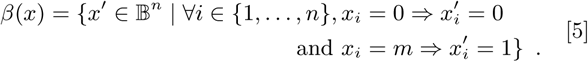

For example with *m* = 2, *β*(012) = {001, 011}.

Then, we say a MN *F* is a refinement of a BN *f* of the same dimension *n* if and only if for every configuration *x* ∈ 𝕄^*n*^, and for every component *i* ∈ {1, …, *n*}, *F*_*i*_(*x*) *<* 0 there exists *x ′* ∈ (*x*) such that *f*_*i*_(*x′*) = 0, and *F*_*i*_(*x*) *>* 0 implies there exists *x ′* ∈ *β* (*x*) such that *f*_*i*_(*x ′*) = 1.

This characterization of BN refinement to MN can be directly extended to ODEs. Indeed, ODEs specify the (real) derivative of the (positive real) value of each component:

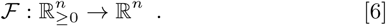

Only the binarization *β* should be adapted in Eq. (5) to reflect that there is no (a priori) upper bounded value *m* for components.

The completeness property states the following. Consider a multivalued refinement *F* of a BN *f* with which there exists an asynchronous trajectory from a multivalued configuration *x* to *y*. Let us write 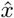 any most-permissive configuration *compatible* with *x*: if *x*_*i*_ = 0, then 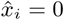, if *x*_*i*_ is the maximum value of *i*, then 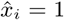, and in the other cases 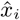 can be either ↗ or ↘. Then, there exists a most-permissive trajectory leading to any of these 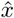 to a most-permissive configuration *ŷ* compatible with *y* and which is consistent with the the changes between *x* and *y*: *ŷ*_*i*_ = ↗ if *y*_*i*_ *> x*_*i*_ and *y*_*i*_ *< m, ŷ*_*i*_ = ↘ if *y*_*i*_ *< x*_*i*_ and *y*_*i*_ *>* 0, and 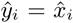 if *y*_*i*_ = *x*_*i*_. As the proof relies solely on the sign of the derivative of the refinement of *f*, the property extends to ODE refinements, which can be seen as multivalued networks with *m* to infinity.

Allowing any state change without restriction would also provide the above guarantee. It appears that if there is a most-permissive trajectory between two binary configurations, then there is a multilevel refinement of the BN showing an asynchronous trajectory between matching multilevel configurations (Theorem 2 in SI.2). Therefore, the completeness property can be achieved only by predicting *at least* the behaviors of MPBNs. In other words, the Most Permissive semantics is the tightest Boolean abstraction of multivalued refinements regarding reachability properties.

### Simpler computational complexity

Most computational analyzes of BNs focus on two elementary dynamical properties: the reachability, which is the existence of a trajectory between two given configurations, and the existence of attractors. Here, we study these properties in term of algorithmic complexity classes. These theoretical results have very concrete implications for the analysis of MPBNs, making the approach scalable to genome-scale networks.

We first recall the bases of computational complexity classes (23): the P class contains algorithms running in time polynomial with the size of its inputs; the NP class contains algorithms running in polynomial time with non-deterministic choices; the PSPACE class contains algorithms running in polynomial *space*. We know that P ⊆ NP ⊆ PSPACE, where “⊆ “can be understood as “simpler”. A problem is complete for a given complexity class if it belongs to and is among the hardest problems of this class. The famous “SAT” problem of determining if a formula expressed in propositional logic (essentially Boolean variables and logic connectors) has a satisfying solution is NP-complete. We do not know if NP = PSPACE, but in practice, NP-complete problems are much more tractable than PSPACE-complete ones by several orders of magnitude. Hereafter, we also refer to the coNP class, which consists of problems for which finding a counter-example belongs to NP, and to the P^NP^ and coNP^coNP^ classes, ordered as follows: coNP ⊆ P^NP^ ⊆ coNP^coNP^ ⊆ PSPACE.

With asynchronous BNs, it is challenging to determine if a trajectory exists between two configurations since, in the worst case, it requires exploring all the possible configurations. With MPBNs, this problem is much simpler thanks to an intriguing property: if there exists a trajectory between two configurations, then there is such a trajectory visiting at most 3*n* configurations. Intuitively, this shortcut corresponds to a particular sequence of state changes: in a first phase, only transitions changing a state from 0 or 1 to ↗ or ↘ take place; In a second phase, only transitions changing states within ↗ and ↘; in a final phase, only transitions changing states from ↗ or ↘ to 1 or 0. Each phase comprises at most *n* transitions, one for each component.

Moreover, finding this shortcut requires exploring at most a quadratic number of transitions in the general case, and only 3*n* whenever the target configuration is in an attractor. The exploration consists of performing as many transitions as possible of the first phase, putting the largest possible number of components in a dynamic state. For each component whose state does not change between the starting and target configuration, it is then necessary to switch the dynamic state back (second phase). If this is not possible, then the exploration is repeated from the beginning while preventing this specific component from changing to a dynamic state (as it would still be impossible to go back to the initial binary state, and the target configuration would not be part of an attractor). Overall, the exploration is thus repeated at most *n* times. Finally, all the transitions of the third phase are applied, which should lead to the target configuration if and only if it is reachable.

On the other hand, determining the possibility of a most-permissive transition is NP-complete in the general case: indeed, the condition “∃ *z* ∈ *γ*(*x*) : *f*_*i*_(*z*) = 1” in Fig. 3 is the SAT problem. For biological networks, it is usual to assume that components cannot have both positive (activator) and negative (inhibitor) direct influences. The resulting BNs are called *locally monotonic*: each local function *f*_*i*_ is monotonic for every component it depends on: increasing the number of activators (resp. inhibitors) of *i* in state 1 can only increase (resp. decrease) the value of *f*_*i*_. Thus, determining the existence of a Boolean interpretation *z* of a most-permissive configuration *x* so that *f*_*i*_(*z*) = 1 comes down to considering activators in dynamic state as 1 and inhibitors in dynamic state as 0, and conversely for *f*_*i*_(*z*) = 0. Therefore, determining the possibility of a most-permissive transition can be done in linear time with locally-monotonic BNs.

The reachability problem in MPBNs can thus be solved in polynomial time whenever *f* is locally monotonic (Theorem 3 in SI.2), a considerable drop in complexity compared to synchronous or asynchronous BNs where the problem is PSPACE-complete. With non-locally monotonic BNs, the reachability problem is in P^NP^.

While the attractors of asynchronous BNs can be complex objects, the attractors of MPBNs are particular mathematical objects called *minimal trap spaces*. A trap space is a hypercube which is closed by *f* : for any vertex *x, f* (*x*) is also a vertex. A trap space is minimal whenever it does not include a different trap space. Attractors in MPBNs have this regular structure because whenever two configurations lying on any diagonal of an hypercube are reachable from each other, they can reach the adjacent configurations as well.

Determining if a configuration *x* ∈ 𝔹^*n*^ belongs to an attractor of *f* is a key problem to identify attractors of a BNs. It is again a PSPACE-complete problem for synchronous and asynchronous BNs. In the case of MPBNs, it boils down to verifying if the trap space containing *x* is minimal, which is at most of complexity coNP for locally monotonic BNs, and at most coNP^coNP^ for non-locally monotonic BNs (Theorem 4 in SI.2). The computation of minimal trap spaces of a BN can be performed efficiently with SAT solvers and related logic programming frameworks (24). On a regular 3.3GHz processor, our implementation of MPBNs can compute reachable attractors of randomly generated scale-free networks (25) with 1,000 components in a fraction of a second, less than 2 seconds with 10,000 components, and less than 50 seconds with 100,000 components (SI.3.C, Fig. S4).

### Validation of MPBNs on actual biological models

An essential feature of logical models is their ability to conclude on the *absence* of certain behaviors. For instance, differentiation processes are modeled using separate attractors representing the final phenotypes and trajectories where configurations are committed to reaching a particular attractor with no possibility to rejoin other differentiation branches. A model allowing any configuration to reach any attractor would indeed be useless without quantitative aspects. We will show that, although enabling more behaviors than (a)synchronous BNs, MPBNs are still constraining and able to capture differentiation and cell fate decisions.

As we have said above, attractors of MPBNs correspond to the minimal trap spaces of the Boolean function. Prior work has shown that these trap spaces match well with the complex attractors of fully asynchronous BNs in many real-world models of biological networks (24). To further assess how MPBNs perform in practice, we reproduced studies on logical models of differentiation using MPBNs instead of fully asynchronous BNs (SI.3).

In the case of a tumor invasion model (8), MPBNs correctly predicted the loss of reachability of apoptotic attractors upon the mutations of p53 and NICD presented in the study (Fig. S2). In the case of T-cell differentiation (9), MPBNs recovered the same reprogramming graph between T-cell types (Fig. S3). After booleanization (26), MPBNs efficiently handled the original large multivalued model of 100 species, whereas the original study had to perform approximations through model reduction. Therefore, the Most Permissive interpretation of BNs is still stringent enough to capture processes that control reachable attractors.

In conclusion, MPBNs are formally guaranteed never to ignore behaviors hidden by artifacts of usual Boolean modeling while still being specific enough to predict differentiation processes, and doing so at a much lower computational cost.

## Discussion

The choice of the dynamical interpretation of BNs has drastic effects on their predictions. Whereas the (fully) asynchronous BN interpretation is often advised for practical applications, it overlooks behaviors emerging from different timescales for the interactions, leading to biases when selecting plausible models. Such misses are due to artifacts of configurations updates. On the contrary, MPBNs offer a framework for reasoning on the qualitative dynamics without making any strong *a priori* hypothesis about the timescale and thresholds of interactions, and without additional parameter.

The state-space explosion triggered by the usual interpretations of BNs is another significant bottleneck for their application in systems biology (3, 27). MPBNs offer drastic gains in computational complexity when analyzing possible trajectories and attractors, both elementary and essential properties, underpining the potential of a model. In practice, the verification of these properties with asynchronous BNs is typically limited to networks with 50 to 100 nodes. On the contrary, deciding the reachability and attractor properties in MPBNs relies on scalable algorithms and does not suffer from the state-space explosion. For the case of locally-monotonic BNs, which is a classical hypothesis for biological networks, the complexity allows addressing very large scale networks, as illustrated in SI.3, with experiments on BNs with up to 100,000 components. Our software tool `mpbn` is available at https://github.com/pauleve/mpbn and integrated in the CoLo-MoTo notebook environment (28).

The prediction of attractors reachable from specific initial conditions, and possibly under various mutant conditions, is at the core of many studies using logical models. While MPBNs can identify the complete set of reachable attractors several orders of magnitude faster than asynchronous BNs, the quantification of the propensities of each attractor, e.g., performed by sampling the trajectories (20, 29), is yet to be explored. In addition to the validation of and the control of predictions from genome-scale models, the complexity breakthrough brought by MPBNs together with their ability to overcome artifacts of Boolean modeling paves the way towards the inference and learning of large-scale logical models from experimental data.

## Acknowledgments

This research was funded by the French Agence Nationale pour la Recherche (ANR) in the context of ANR-FNR project “AlgoReCell” (ANR-16-CE12-0034)

## Authors’ contributions

LP, TC, SH designed the research; LP defined MPBNs and demonstrated Theorems 1,3,4, implemented code, performed experiments, and initially drafted the manuscript; JK demonstrated Theorem 2; LP, JK, TC, SH wrote the manuscript.

