## Supplementary Information for "Reconciling Qualitative, Abstract, and Scalable Modeling of Biological Networks"

**4 I3-FFL network motif** **12**

#### List of Figures

- S1 Incoherent feed-forward loop of type 3 (a) and its associated Boolean logic for nodes activities (b). Whereas theoretical and experimental studies show a possible activation of the output (c), usual Boolean networks analysis cannot predict this transient behavior (d). Conversely, the Most Permissive interpretation can reproduce this behavior, (e) showing one of its possible execution. A configuration is represented by a pile of 3 squares, where the top square represents the state of the first component, and so forth. A white square represent the inactive (0) state; a blue square represents the active (1) state; the increasing state $\nearrow$ is represented by a top-right white and bottom-left blue square, and decreasing state $\searrow$ by a top-right blue and bottom-left white square. . . . 13
- S2 Influence graph of the tumour invasion model by Cohen et al. (2015) . . . 14
- S3 Influence graph of the T-cell differentiation model by Abou-Jaoudé et al. (2015) . . . 15
- S4 Computation times on a 3GHz CPU obtained using `mpbn` software tool on BNs generated with random scale-free influence graph for the computation of a single attractor, the enumeration of 1,000 attractor, and the computation of attractors reachable from random initial conditions. See notebook "Scalability on large random BNs.ipynb" available at doi:10.5281/zenodo.3936123. . . . 16

### Supporting Information Text

### 1. Formal Background on Boolean networks

**Notations** The Boolean domain is denoted by $\mathbb{B} := \{0,1\}$. Given a *configuration* $x \in \mathbb{B}^n$ and $i \in [n]$, we denote $x_i$ the $i^{\text{th}}$ component of $x$, so that $x = x_1 \dots x_n$, and $\bar{x}$ the complement of $x$, i.e., $\forall i \in [n]$, $\bar{x}_i = 1 - x_i$. Given two configurations $x, y \in \mathbb{B}^n$, the components having a different state are noted $\Delta(x,y) := \{i \in [n] \mid x_i \neq y_i\}$. Symbol $\wedge$ denotes the logical conjunction, $\vee$ the disjunction, and $\neg$ the negation. Given a finite set $S$, $|S|$ is its cardinality.

#### A. Boolean networks.

**Definition 1.1** (Boolean network)**.** A Boolean network (BN) of dimension $n$ is a function $f : \mathbb{B}^n \to \mathbb{B}^n$. For each $i \in [n]$, $f_i : \mathbb{B}^n \to \mathbb{B}$ denotes the local function of its $i$th component.

**Definition 1.2** (Locally-monotonic BN)**.** A BN $f : \mathbb{B}^n \to \mathbb{B}^n$ is *locally monotonic* whenever for each component $i \in \{1, \dots, n\}$, there exists an ordering of components $\preceq^i \in \{\leq, \geq\}^n$ such that $\forall x, y \in \mathbb{B}^n, (x_1 \preceq^i_1 y_1 \wedge \dots \wedge x_n \preceq^i_n y_n) \Rightarrow f_i(x) \leq f_i(y)$.

**Example 1.** *The BN $f$ of dimension 3 defined as*

$$f_1(x) = x_3 \wedge (\neg x_1 \vee \neg x_2)$$
$$f_2(x) = x_3 \wedge x_1$$
$$f_3(x) = x_1 \vee x_2 \vee x_3 \ ,$$

*is locally monotonic, for instance with $\preceq^1 = (\geq, \geq, \leq)$ and $\preceq^2 = \preceq^3 = (\leq, \leq, \leq)$.*

#### B. Update semantics.

BN semantics are expressed as irreflexive binary relations between the configurations. We use the symbol $\to$ decorated with the Boolean function and a symbol representing the semantics.

**Definition 1.3** (Synchronous semantics)**.**

$$\forall x, y \in \mathbb{B}^n \quad x \xrightarrow[\text{s}]{f} y \overset{\Delta}{\iff} x \neq y \wedge y = f(x) \ .$$

**Definition 1.4** (Fully asynchronous semantics)**.**

$$\forall x, y \in \mathbb{B}^n, \quad x \xrightarrow[\text{a1}]{f} y \overset{\Delta}{\iff} \exists i \in [n] : \Delta(x,y) = \{i\} \wedge y_i = f_i(x) \ .$$

**Definition 1.5** (Asynchronous semantics)**.**

$$\forall x, y \in \mathbb{B}^n, \quad x \xrightarrow[\text{a}]{f} y \overset{\Delta}{\iff} x \neq y \wedge \forall i \in \Delta(x,y), y_i = f_i(x) \ .$$

Given a semantics $\sigma$, we write $\xrightarrow[\sigma]{f}{}^*$ the reflexive and transitive closure of the binary relation $\xrightarrow[\sigma]{f}$. Thus, $x \xrightarrow[\sigma]{f}{}^* y$ if and only if $x = y$ or there exists a sequence $x \xrightarrow[\sigma]{f} x' \xrightarrow[\sigma]{f} \cdots \xrightarrow[\sigma]{f} y$. The set of configurations which are in such a relation with a configuration $x$ is given by $\rho^f_\sigma(x)$:

$$\rho^f_\sigma(x) := \{y \in \mathbb{B}^n \mid x \xrightarrow[\sigma]{f}{}^* y\} \ . \qquad [1]$$

#### C. Dynamical properties.

**Definition 1.6** (Fixed point)**.** A configuration $x \in \mathbb{B}^n$ is a *fixed point* of the BN $f$ with semantics $\sigma$ whenever

$$\rho^f_\sigma(x) = \{x\} \ .$$

**Definition 1.7** (Reachability)**.** Given two configurations $x, y \in \mathbb{B}^n$ of a BN $f$ with semantics $\sigma$, $y$ is *reachable* from $x$ whenever

$$y \in \rho^f_\sigma(x) \ .$$

**Definition 1.8** (Attractor)**.** A non-empty set of configurations $A \subseteq \mathbb{B}^n$ is an *attractor* of the BN $f$ with semantics $\sigma$ whenever

$$\forall x \in A, \quad \rho^f_\sigma(x) = A \ .$$

#### D. Computational Complexity of Dynamical Properties with (A)synchronous Semantics.

Let us fix a BN $f$ of dimension $n$.

***D.1. Fixed points.*** Remark that with synchronous, fully asynchronous, and asynchronous semantics, $x \in \mathbb{B}^n$ is a fixed point if and only if $f(x) = x$.

**Proposition 1.** *Deciding if there exists $x \in \mathbb{B}^n$ such that $f(x) = x$ is NP-complete.*

*Proof.* By reduction of the SAT problem (1). □

#### D.2. Reachability.

**Proposition 2.** *Given two configurations $x, y \in \mathbb{B}^n$, deciding if $y \in \rho^f_\sigma(x)$ with $\sigma \in \{s, a1, a\}$ is PSPACE-complete.*

*Proof.* As there is at most $2^n$ configurations to explore, the problem is at most in PSPACE, as it sufficient to apply non-deterministically at most $2^n - 1$ transitions from $x$ using a counter on $n$ bits.

With the synchronous semantics, the PSPACE-hardness derives by reduction of the reachability problem in reaction systems, a subclass of synchronous BNs (2).

With fully asynchronous and asynchronous semantics, the PSPACE-hardness derives by reduction of the reachability problem in synchronous BNs. Indeed, similarly to cellular automata (3), one can define a BN $f'$ so that asynchronous and fully asynchronous semantics give reachability relations that are equivalent with the synchronous semantics of $f$.

A possible construction is to decompose a synchronous transition in several steps which can be performed asynchronously. This involves 3 stages: (a) the computation of the next value for each component $i \in [n]$; (b) the application of the new state for each component; (c) the reset of components introduced by the construction. We give here an encoding as a BN $f'$ with $3n+2$ dimensions: one component z for which the state 1 triggers the reset of additional components (except z); one component w for which the state 0, assuming z has state 0, triggers the computation stage (a), and the state 1 triggers the application stage (b). For each component $i \in [n]$ of $f$, two components $\mathrm{c}i$ and $\bar{\mathrm{c}}i$ are defined, for which the state 1 specify respectively if $f_i$ is true or false. The end of computation stage (a) is detected whenever for each component $i \in [n]$, either $\mathrm{c}i$ or $\bar{\mathrm{c}}i$ are in state 1. Component w then switch to state 1; then, components $i$ switch to state 0 if and only if $\bar{\mathrm{c}}i$ is 1 and to state 1 if and only if $\mathrm{c}i$ is 1. The end of application stage (b) is detected whenever all the components $i \in [n]$ have been updated. Component z then witch to state 1 which will trigger the switch to state 0 of components w, $\mathrm{c}i$ and $\bar{\mathrm{c}}i$. Finally, component $z$ switch back to state 0, which allows components $\mathrm{c}i$ and $\bar{\mathrm{c}}i$ computing the next state of each components $i \in [n]$. This network $f' : \mathbb{B}^{3n+2} \to \mathbb{B}^{3n+2}$ can be formally defined as follows, where $x_{1..n}$ denotes the configuration $x$ truncated at the first $n$ components:

$$
\begin{aligned}
f'_i(x') &= ((\neg x'_{\mathrm{w}} \vee x'_{\mathrm{z}}) \wedge x'_i) \vee (x'_{\mathrm{w}} \wedge \neg x'_{\mathrm{z}} \wedge x'_{\mathrm{c}i}) \\
f'_{\mathrm{c}i}(x') &= \neg x'_{\mathrm{z}} \wedge \big((\neg x'_{\mathrm{w}} \wedge f_i(x'_{1..n})) \vee (x'_{\mathrm{w}} \wedge x'_{\mathrm{c}i})\big) \\
f'_{\bar{\mathrm{c}}i}(x') &= \neg x'_{\mathrm{z}} \wedge \big((\neg x'_{\mathrm{w}} \wedge \neg f_i(x'_{1..n})) \vee (x'_{\mathrm{w}} \wedge x'_{\bar{\mathrm{c}}i})\big) \\
f'_{\mathrm{w}}(x') &= \neg x'_{\mathrm{z}} \wedge \Big(\Big(x'_{\mathrm{w}} \vee \textstyle\bigwedge_{i\in[n]}(x'_{\mathrm{c}i} \vee x'_{\bar{\mathrm{c}}i})\Big)\Big) \\
f'_{\mathrm{z}}(x') &= \Big(x'_{\mathrm{w}} \wedge \textstyle\bigwedge_{i\in[n]}(x'_{\mathrm{c}i} \Leftrightarrow x'_i \wedge x'_{\bar{\mathrm{c}}i} \Leftrightarrow \neg x'_i)\Big) \vee \Big(x'_{\mathrm{z}} \wedge \Big(x'_{\mathrm{w}} \vee \textstyle\bigvee_{i\in[n]}(x'_{\mathrm{c}i} \vee x'_{\bar{\mathrm{c}}i})\Big)\Big)
\end{aligned}
$$

It results that for all pairs of configurations $x, y \in \mathbb{B}^n$,

$$
y \in \rho^f_{\mathrm{s}}(x) \iff y0^{2n+2} \in \rho^{f'}_{\mathrm{a}}(x0^{2n+2}) \iff y0^{2n+2} \in \rho^{f'}_{\mathrm{a1}}(x0^{2n+2})
$$

where $0^{2n+2}$ is the 0 vector of dimension $2n+2$ and $y0^{2n+2}$ denotes its concatenation to $y$. ☐

#### D.3. Attractors.

**Proposition 3.** *Given a configuration $x \in \mathbb{B}^n$, deciding if $x$ belongs to an attractor of $f$ with semantics $\sigma \in \{s, a1, a\}$ is PSPACE-complete.*

*Proof.* The problem is in PSPACE as one can solve it through its complementary: $x$ does not belong to any attractor if and only if there exists a configuration $y$ such that $y$ is reachable from $x$ and $x$ is not reachable from $y$.

For the synchronous semantics, the PSPACE-hardness derives by reduction of the same problem in synchronous reaction systems, a subclass of BNs (2). Then, the proof can be lifted to asynchronous and fully asynchronous semantics by the reduction of the problem with the synchronous semantics (e.g., by using the construction in the previous section). ☐

### 2. Most Permissive Boolean Networks

**A. Formal definition of the most permissive semantics.** We give two different definitions which are equivalent in term of reachability properties. The first one introduces dynamic states, the second one relies on the computation of hypercubes.

***A.1. With dynamic states.*** A most-permissive configuration assigns to each BN component one state among four, noted $\mathbb{P} := \{0, \nearrow, \searrow, 1\}$. The possible binary interpretations of a configuration $x \in \mathbb{P}^n$ are denoted by

$$\gamma(x) := \{\tilde{x} \in \mathbb{B}^n \mid \forall i \in [n], x_i \in \mathbb{B} \Rightarrow \tilde{x}_i = x_i\} \ . \tag{2}$$

The semantics is defined as an irreflexive binary relation between configurations in $\mathbb{P}^n$:

**Definition 2.1** (Most permissive semantics $\xrightarrow[\text{mp}]{f}$)**.**

$$\forall x, y \in \mathbb{P}^n, \quad x \xrightarrow[\text{mp}]{f} y \overset{\Delta}{\iff} \exists i \in [n] : \Delta(x, y) = \{i\}$$

$$\wedge\, y_i = \begin{cases} \nearrow & \text{if } x_i \neq 1 \wedge \exists \tilde{x} \in \gamma(x) : f_i(\tilde{x}) \\ 1 & \text{if } x_i = \nearrow \\ \searrow & \text{if } x_i \neq 0 \wedge \exists \tilde{x} \in \gamma(x) : \neg f_i(\tilde{x}) \\ 0 & \text{if } x_i = \searrow \end{cases}$$

The set of binary configurations reachable from $x \in \mathbb{B}^n$ with the most permissive semantics is given by

$$\rho^f_{\text{mp}}(x) := \{y \in \mathbb{B}^n \mid x \xrightarrow[\text{mp}]{f}{}^* y\} \ . \tag{3}$$

The following figure shows the automaton of the state change of a component $i$ in the most permissive semantics, following notations of Def. 2.1. The labels $f_i(\tilde{x})$ and $\neg f_i(\tilde{x})$ on edges are the conditions for firing the transitions, where $\tilde{x} \in \gamma(x)$; the label $\epsilon$ indicates transitions that can be done without condition:

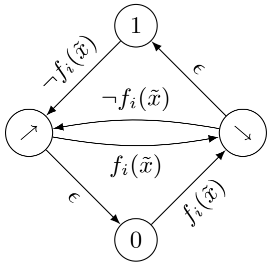

With the given definition, only one automaton is updated at a time. However, it is equivalent to allow any number of simultaneous changes, as long as fully asynchronous updates are considered.

Given a configuration $x \in \mathbb{P}^n$, one can remark that as long as only transitions towards dynamic states $\nearrow$ or $\searrow$ are performed, then the set of binary interpretations $\gamma$ is growing. As a consequence, the ordering of such transitions does not matter.

**Proposition 4.** *Given a BN $f$ of dimension $n$, $\forall x, y \in \mathbb{P}^n$ such that $x \xrightarrow[\text{mp}]{f} y$ and $\forall j \in \Delta(x, y) : y_j \notin \mathbb{B}$, $\gamma(x) \subseteq \gamma(y)$.*

Given a configuration $x \in \mathbb{B}^n$, if we consider any reachable configuration where changed components are in an dynamic state, and from which there is no more transitions from binary states towards dynamic states, then the set of binary interpretation of this later configuration includes the set of all binary configurations reachable from $x$:

**Proposition 5.** *Given a BN $f$ of dimension $n$ and a binary configuration $x \in \mathbb{B}^n$, let us consider a configuration $z \in \mathbb{P}^n$ such that $x \xrightarrow[\text{mp}]{f}{}^* z$, $\forall i \in \Delta(x, z) : z_i \notin \mathbb{B}$, and there is no $z' \in \mathbb{P}^n$ such that $z \xrightarrow[\text{mp}]{f} z$ with for $j \in \Delta(z, z')$, $z_j \in \mathbb{B}$ and $z'_j \notin \mathbb{B}$, then $\rho^f_{\text{mp}}(x) \subseteq \gamma(z)$.*

***A.2. With hypercubes.*** The dynamic states might suggest that the most permissive semantics is close to multivalued networks with 4 states. However, notice that states $\mathbb{P}$ are not totally ordered by the transitions, as it is required by multivalued networks.

We give here an equivalent definition of $\rho^f_{\text{mp}}$ which does not relies on these dynamic states, but on the computation of hypercubes closed by $f$. An hypercube within $\mathbb{B}^n$ has a set of components being fixed to a Boolean state, and the others being free (noted with $*$).

**Definition 2.2** (Hypercube)**.** An *hypercube* $h$ of dimension $n$ is a vector in $(\mathbb{B} \cup \{*\})^n$. The set of its associated configurations is denoted by $c(h) := \{x \in \mathbb{B}^n \mid \forall i \in [n], h_i \neq * \Rightarrow x_i = h_i\}$.

Given two hypercubes $h, h' \in (\mathbb{B} \cup \{*\})^n$, $h$ is *smaller* than $h'$ if and only if $\forall i \in [n], h'_i \neq * \Rightarrow h_i = h'_i$. An hypercube is *minimal* if there is no different hypercubes smaller than it.

An hypercube $h$ is *closed* by $f$ whenever for each configuration $x \in c(h)$, $f(x) \in c(h)$.

An hypercube closed by $f$ is also known as a *trap space*; if it is minimal, it is a *minimal trap space*.
We generalize the notion of closure by allowing restricting the set of components which should be closed.

**Definition 2.3** ($K$-closed hypercube)**.** Given a subset of components $K \subseteq [n]$, an hypercube $h \in (\mathbb{B} \cup \{*\})^n$ is $K$-closed by $f$ whenever for each configuration $x \in c(h)$, for each component $i \in K$, $h_i \in \{*, f_i(x)\}$.

Remark: an hypercube is closed if and only if it is $[n]$-closed.

**Example 2.** *Let us consider the BN $f : \mathbb{B}^3 \to \mathbb{B}^3$ with $f_1(x) := \neg x_2$, $f_2(x) := \neg x_1$, et $f_3(x) := \neg x_1 \wedge x_2$. The hypercube $01*$ is closed by $f$, with $c(01*) = \{010, 011\}$. The hypercube $1{*}0$ is the smallest hypercube $\{2,3\}$-closed by $f$ containing $110$; it is not closed by $f$, nor the smallest hypercube $\{2,3\}$-closed by $f$ containing $100$.*

Starting from a binary configuration $x \in \mathbb{B}^n$, the most permissive semantics can be expressed using the computation of smallest hypercubes containing $x$ and which are $K$-closed by $f$, for every $K$:

- $x$ is the unique hypercube $\emptyset$-closed by $f$ containing $x$;
- the change of state of component $i \in [n]$ to $\nearrow$ or $\searrow$ produces a configuration $x'$ where $\gamma(x')$ correspond to the hypercube $h \in (\mathbb{B} \cup \{*\})^n$ with $h_i = *$ and for each other component $j \in [n], j \neq i$, $h_j = x_j$. Thus, $h$ is the smallest hypercube $\{i\}$-closed by $f$ and containing $x$;
- by considering only the change of states towards $\nearrow$ and $\searrow$, the most permissive semantics progressively enlarges the hypercubes along the modified components, and each step results in a smallest hypercube $K$-closed by $f$ and containing $x$, for every $K \subseteq [n]$.

With the most permissive semantics, the change of state of a component from a dynamic to a Boolean state is without condition, and is solely determined by its current dynamic state: 1 from $\nearrow$ and 0 from $\searrow$. Thus, starting from an initial configuration which is binary, a component can be in the state $\nearrow$ only if a preceding configuration $x' \in \mathbb{P}^n$ was such that $\exists z \in \gamma(x')$ with $f_i(z) = 1$ (resp. $\searrow$ if $f_i(z) = 0$).
The following proposition establishes the correspondence with the initial definition with dynamic states:

**Proposition 6.** *Given a BN $f$ of dimension $n$ and two configurations $x, y \in \mathbb{B}^n$, $y \in \rho^f_{\mathrm{mp}}(y)$ if and only if there exists $K \subseteq [n]$ such that the smallest $K$-closed hypercube $h$ and containing $x$ verifies (1) $y \in c(h)$, and (2) $\forall i \in K$, there exists a configuration $z \in c(h)$ such that $f_i(z) = y_i$.*

**Example 3.** *Here below are examples of smallest $K$-closed hypercubes containing la configuration $000$ (left), $010$ (top right), and $011$ (bottom right) for the BN $f$ of dimension $3$ defined by $f_1(x) := \neg x_2$, $f_2(x) := \neg x_1$, $f_3(x) := \neg x_1 \wedge x_2$. Configurations belonging to the hypercube are highlighted in bold; these verifying the reachability property are boxed. The hypercube $011$ is only one which is closed by $f$ and minimal.*

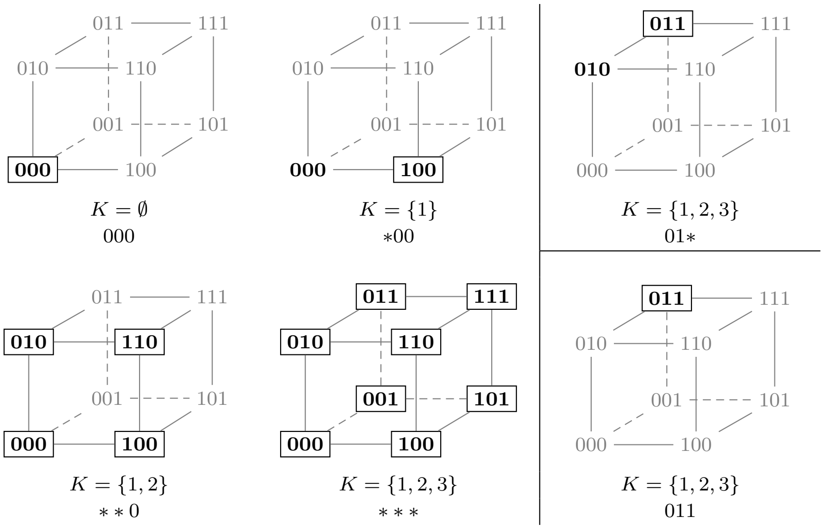

**B. Relation with quantitative refinements.** Multivalued networks (MNs) are a generalization of BNs where the components can take values in a finite discrete domain. Let us denote the possible values as $\mathbb{M} := \{0, 1, \ldots, m\}$ for some integer $m$. Without loss of generality, we assume the same domain of values for all the components.

**Definition 2.4** (Multivalued network)**.** A *multivalued network* (MN) of dimension $n$ over a value range $\mathbb{M} = \{0, 1, \ldots, m\}$ is a function $F : \mathbb{M}^n \to \{-1, 0, 1\}^n$.

A configuration of a MN of dimension $n$ is a vector $x \in \mathbb{M}^n$. Given two configurations $x, y \in \mathbb{M}^n$, the components that differ are noted $\Delta(x, y) := \{i \in [n] \mid x_i \neq y_i\}$.

**Definition 2.5** (Asynchronous semantics)**.** Given a multivalued network $F$, the binary irreflexive relation $\xrightarrow[\text{a}]{F} \subseteq \mathbb{M}^n \times \mathbb{M}^n$ is defined as:

$$x \xrightarrow[\text{a}]{F} y \overset{\Delta}{\iff} \forall i \in \Delta(x,y), y_i = x_i + F_i(x) \enspace .$$

We write $\xrightarrow[\text{a}]{F}^*$ for the transitive closure of $\xrightarrow[\text{a}]{F}$.

We now define a notion of *multivalued refinement* of a BN, which formalizes the intuition that the value changes defined by the multivalued network are compatible with those of the BN. The refinement criteria relies on a *binarization* of the multivalued configuration. An appropriate binarization necessarily quantifies 0 as Boolean 0 and $m$ as 1, and is free for the other dynamic states. Let us denote by $\beta(x)$ the set of possible binarization of configuration $x \in \mathbb{M}^n$:

$$\beta(x) := \{x' \in \mathbb{B}^n \mid \forall i \in [n], x_i = 0 \Rightarrow x'_i = 0 \wedge x_i = m \Rightarrow x'_i = 1\} \enspace . \qquad [4]$$

**Definition 2.6** (Multivalued refinement)**.** A multivalued network $F$ of dimension $n$ over a value range $\mathbb{M}$ *refines* a BN $f$ of equal dimension $n$ if and only if for every configuration $x \in \mathbb{M}^n$ and every $i \in [n]$,

$$F_i(x) > 0 \Rightarrow \exists x' \in \beta(x) : f_i(x') = 1 \wedge F_i(x) < 0 \Rightarrow \exists x' \in \beta(x) : f_i(x') = 0 \enspace .$$

This characterization of refinement can be readily extended to ODEs: similarly to multivalued networks, ODEs specify the derivative of the (positive) real value of each component along the continuous time $t$:

$$\frac{d\mathbb{F}(t,x)}{dt} = \mathcal{F}(x) \qquad \text{with } \mathcal{F} : \mathbb{R}^n_{\geq 0} \to \mathbb{R}^n \enspace . \qquad [5]$$

Here, $\mathcal{F}(x)$ is the derivative of $\mathbb{F}(t,x)$ along time $t$ in function of continuous configurations $x$; $\mathbb{F}$ being usually unknown. ODEs can be seen thus be seen as MNs with $m$ going to infinity and with synchronous semantics: $\mathcal{F}$ model the simultaneous evolution of all the components.

The admissible binarizations $\beta$ should be slightly adapted to reflect the absence *a priori* of maximum value: $\beta(x) := \{x' \in \mathbb{B}^n \mid \forall i \in [n], x_i = 0 \Rightarrow x'_i = 0\}$. Then, the definition of refinement is identical.

***B.1. Theorem 1 (Completeness).*** Let us consider a BN $f$ of dimension $n$ and any multivalued refinement $F$ with $m$ values. A *most-permissive interpretation* of a multivalued configuration is a configuration in $\mathbb{P}^n$ where components having extreme states in the multivalued configuration have the corresponding extreme states in the most permissive configuration, and otherwise are either $\nearrow$ or $\searrow$. Let us denote these interpretations by

$$\alpha(x) := \{\hat{x} \in \mathbb{P}^n \mid x_i = 0 \Leftrightarrow \hat{x}_i = 0 \wedge x_i = m \Leftrightarrow \hat{x}_i = m\} \qquad [6]$$

Then, Theorem 1 states that for any asynchronous transition from $x$ to $y$ ($x \xrightarrow[\text{a}]{F} y$), there is a most permissive trajectory from any corresponding most permissive configuration $\hat{x} \in \alpha(x)$ to a configuration $\hat{y} \in \alpha(y)$ where the state of each component is consistent with the changes between $x$ and $y$.

**Theorem 1.** *Given a BN $f$ of dimension $n$, for any multivalued network $F : \mathbb{M}^n \to \{-1,0,1\}^n$ being a refinement of $f$,*

$$\forall x,y \in \mathbb{M}^n, \quad x \xrightarrow[\text{a}]{F} y \Longrightarrow \forall \hat{x} \in \alpha(x), \exists \hat{y} \in \alpha(x) : \hat{x} \xrightarrow[\text{mp}]{f}^* \hat{y} \textit{ with } \forall i \in [n], \hat{y}_i = \begin{cases} \nearrow & \textit{if } y_i > x_i \wedge y_i < m \\ \searrow & \textit{if } y_i < x_i \wedge y_i > 0 \\ 0 & \textit{if } y_i = 0 \\ 1 & \textit{if } y_i = m \\ \hat{x}_i & \textit{otherwise.} \end{cases}$$

*Proof.* From MN semantics, for each component $i \in \Delta(x,y)$, whenever $y_i > x_i$ (resp. $y_i < x_i$), necessarily $F_i(x) > 0$ (resp. $F_i(x) < 0$). From the refinement property, there exists a binarization $x' \in \beta(x)$ such that $f_i(x) = 1$ (resp. $f_i(x) = 0$). Now remark that for any $\hat{x} \in \alpha(x)$, $x' \in \beta(\hat{x})$. Therefore, for each component $i \in \Delta(x,y)$, if $y_i > x_i$ and $\hat{x}_i \neq \nearrow$, the $i$ can change to state $\nearrow$, and if $y_i < x_i$ and $\hat{x}_i \neq \searrow$, the $i$ can change to state $\searrow$. By Proposition 4, these transitions can be applied in any order; let us denote by $z$ the obtained configuration. Finally, for each component $i \in \Delta(x,y)$ where $y_i = 0$ (resp. $y_i = m$), remark that $z_i = \searrow$ (resp. $z_i = \nearrow$), thus it can change to state 0 (resp. 1), in any order. Therefore, $\hat{x} \xrightarrow[\text{mp}]{f}^* \hat{y}$. ☐

Remark that the theorem considers asynchronous transition, which includes any restrictions (synchronous, fully asynchronous, sequential, ...).

As the proof relies solely on the sign of the derivative of the refinement of $f$, the property extends to ODE refinements, which can be seen as MN with $m$ to infinity and with synchronous semantics. The function $\alpha$ then becomes

$$\alpha(x) := \{\hat{x} \in \mathbb{P}^n \mid \forall i \in [n], x_i = 0 \Leftrightarrow \hat{x}_i = 0 \wedge \hat{x}_i \neq 1\} \qquad [7]$$

**Corollary 1.** *For any ODE system* $\mathcal{F} : \mathbb{R}^n_{\geq 0} \to \mathbb{R}^n$ *refining a BN* $f$ *of dimension* $n$*,*

$$\forall x \in \mathbb{R}^n_{\geq 0}, \forall \hat{x} \in \alpha(x), \quad \hat{x} \xrightarrow[\text{mp}]{f}{}^* \hat{y} \quad \text{with } \forall i \in [n], \hat{y}_i = \begin{cases} \nearrow & \text{if } F_i(x) > 0 \\ \searrow & \text{if } F_i(x) < 0 \wedge x_i > 0 \\ \hat{x}_i & \text{otherwise.} \end{cases}$$

Remark that a BN $f$ is a multivalued refinement of itself with $\mathbb{M} = \mathbb{B}$ and for each $i \in [n]$, $F_i(x) = 1$ if $f_i(x), -1$ otherwise. Therefore another corollary of the above theorem is that the most permissive semantics of BNs simulates the asynchronous semantics of $f$:

**Corollary 2.** *Given a BN* $f$ *of dimension* $n$*,*

$$\forall x, y \in \mathbb{B}^n, \quad x \xrightarrow[\text{a}]{f}{}^* y \Longrightarrow x \xrightarrow[\text{mp}]{f}{}^* y \enspace .$$

Thus, the number of attractors with the most permissive semantics is at most the number of attractor with update semantics.

***B.2. Theorem 2 (Minimality).*** Whereas complete, one should wonder whether the most permissive semantics introduce spurious behaviors. We prove in this section that the most permissive semantics is the tightest abstraction of multivalued refinements with respect to reachability properties.

First, Proposition 7 ensures that if there exists a most-permissive trajectory between two Boolean configurations $x$ and $y \in \mathbb{B}^n$, then there exists a multilevel refinement of the BN which allows an asynchronous trajectory between corresponding configurations $m.x$ and $m.y$ with $m = 2$. The idea is to construct a MN which can reproduce the shortcut trajectory, with dynamic states identified to an intermediate state 1 of the MN: in a first phase, components increase to 1 (possibly fully-asynchronously), then a last synchronous step leads to the target $2.y$ configuration.

Then, we introduce the notion of trace refinement witch matches most permissive trajectories with MN asynchronous trajectories having coherent successions of states, both with respect to admissible most-permissive interpretation, and with respect to derivatives: whenever a component $i$ changes to the dynamic state $\nearrow$ (resp. $\searrow$), $F_i$ is positive (resp. negative) in the corresponding multivalued configuration. Theorem 2 establishes for any most permissive trajectory, there exists a MN refinement with $m = 3$ which admits a matching asynchronous trajectory.

Therefore, the most permissive semantics introduces no spurious behavior with respect to the admissible refinements of a BN $f$.

**Proposition 7.** *For any BN* $f$ *of dimension* $n$ *and any pair of configurations* $x, y \in \mathbb{B}^n$*, if* $y$ *is reachable from* $x$ *with the most permissive semantics, then there exists a MN* $F$ *with* $m$ *values which is a refinement of* $f$ *and where* $m.y$ *is reachable from* $m.x$ *with the asynchronous semantics.*

*Proof.* Let $K \subseteq [n]$ be the smallest subset of components verifying Proposition 6. We now define a sequence of configurations $x, x', \dots, x^{(|K|)} \in \mathbb{P}^n$ to be arbitrary such that $\forall 0 < i \leq |K|$, $x^{(i-1)} \xrightarrow[\text{mp}]{f} x^{(i)}$ and $j \in \Delta(x^{(i-1)}, x^{(i)}) \Longrightarrow j \in K \wedge x_j^{(i-1)} \in \mathbb{B} \wedge x_j^{(i)} \notin \mathbb{B}$. Note that such a sequence is guaranteed to exist thanks to $K$ being minimal.

We define another sequence of configurations $z, z', \dots, z^{(|K|)} \in \{0,1,2\}^n$ as the multivalued equivalent of $x, x', \dots, x^{(|K|)}$: $\forall 0 \leq i \leq |K|$ and $\forall j \in [n]$, $x_j^{(i)} \in \mathbb{B} \Longrightarrow z_j^{(i)} = 2.x_j^{(i)}$ and $x_j^{(i)} \notin \mathbb{B} \Longrightarrow z_j^{(i)} = 1$.

We now construct the coveted MN $F$ with 3 values. based on $z, z', \dots, z^{(|K|)}$ as follows:

- For any $0 \leq i < |K|$, $F(z^{(i)}) = z^{(i+1)} - z^{(i)}$.
- $F(z^{|K|}) = 2.y - z^{|K|}$. ($2.y - z^{|K|} \in \{-1,0,1\}^n$ thanks to $y$ being in the smallest $K$-closed hypercube containing $x$.)
- For any other $z \in \{0,1,2\}^n$, $F(z) = 0^n$.

Clearly, $2.y$ is reachable from $2.x = z$ in $F$ with synchronous semantics. What remains to be proven is that $F$ is a refinement of $f$. Nothing needs to be shown for cases when $F$ returns 0, let thus first $0 \leq i < |K|$ and $\{j\} = \Delta(z^{(i)}, z^{(i+1)})$. Let us further assume $F_j(z^{(i)}) = 1$ as the $F_j(z^{(i)}) = -1$ case is symmetric. We need to show $\exists \tilde{x} \in \beta(z^{(i)})$ such that $f(\tilde{x})$. By definition, $x^{(i+1)} = \nearrow$, thus $\exists \tilde{x} \in \gamma(x^{(i)})$ such that $f(\tilde{x})$. Since for any $j \in [n]$, $z_j^{(i)} = 1$ exactly when $x_j^{(i)} \notin \mathbb{B}$, we have $\gamma(x^{(i)}) \subseteq \beta(z^{(i)})$.

Finally, let us consider $z^{|K|}$. We need to show $\forall j \in \Delta(z^{(|K|)}, 2.y)$, $\exists \tilde{x} \in \beta(z^{(|K|)})$, $f(\tilde{x}) = y_j$. By definition, we have $\Delta(z^{(|K|)}, 2.y) = K$. Since $K$ verifies Property 6, we know $\forall j \in K$, $\exists \tilde{x} \in c(h)$, $f(\tilde{x}) = y_j$, where $h$ is the smallest $K$-closed hypercube containing $x$. By definition of $x^{(|K|)}$, $\forall j \in K$, $x_j^{(|K|)} \notin \mathbb{B}$ and thus $c(h) \subseteq \gamma(x^{(|K|)})$. Furthermore, since for any $j \in [n]$, $z_j^{(|K|)} = 1$ exactly when $x_j^{(|K|)} \notin \mathbb{B}$, we have $\gamma(x^{(|K|)}) \subseteq \beta(z^{(|K|)})$.

□

**Definition 2.7** (Trace Refinement)**.** Given a BN $f$ of dimension $n$ and a multivalued refinement $F : \mathbb{M}^n \to \{-1,0,1\}^n$ of $f$. Let $x, x', \dots, x^{(k)} \in \mathbb{P}^n$ be a finite sequence of configurations such that $\forall 0 < i \leq k$, $x^{(i-1)} \xrightarrow[\text{mp}]{f} x^{(i)}$ (finite trace of $f$ with the most permissive semantics).

Then a finite sequence $y, y', \ldots, y^{(l)} \in \mathbb{M}^n$ such that $\forall 0 < i \leq l$, $y^{(i-1)} \xrightarrow[\text{a}]{F} y^{(i)}$, is a trace refinement of $x, x', \ldots, x^{(k)}$ if there exists a function $\kappa : \{0, \ldots, k\} \to \{0, \ldots, l\}$ (trace refinement function) satisfying the following requirements:

1. $\kappa$ is non-decreasing, i.e. $i < j \implies \kappa(i) \leq \kappa(j)$;
2. $\kappa(0) = 0$ and $\kappa(k) = l$;
3. $\forall j \in [n]$, $x_j = y_j$ and for each $0 < i \leq k$, $(x_j^{(i)} = 0 \implies y_j^{(\kappa(i))} < m) \wedge (x_j^{(i)} = 1 \implies y_j^{(\kappa(i))} > 0)$;
4. For each $0 < i \leq k$ such that $x_j^{(i)} \notin \mathbb{B}$ where $\{j\} = \Delta(x^{(i-1)}, x^{(i)})$, $x_j^{(i)} = \nearrow \implies F_j(y^{(\kappa(i-1))}) = 1$ and $x_j^{(i)} = \searrow \implies F_j(y^{(\kappa(i-1))}) = -1$.

**Theorem 2.** *For any BN $f$ of dimension $n$ and any sequence of configurations $x, x', \ldots, x^{(k)} \in \mathbb{P}^n$ such that $x \in \mathbb{B}^n$ and $\forall 0 < i \leq k$, $x^{(i-1)} \xrightarrow[\text{mp}]{f} x^{(i)}$, there exists a MN $F : \mathbb{M}^n \to \{-1, 0, 1\}^n$ which is a refinement of $f$ and has a trace refinement $y, y', \ldots, y^{(l)} \in \mathbb{M}^n$ of $x, x', \ldots, x^{(k)}$.*

*Proof.* We construct $F$ and $y, y', \ldots, y^{(l)}$ iteratively along the sequence $x, x', \ldots, x^{(k)}$. For each step $i \in \{0, \ldots, k\}$ we maintain that the constructed network $F$ is a refinement of of $f$ and $y, y', \ldots, y^{(l_i)}$ is a trace refinement of $x, x', \ldots, x^{(i)}$.

Let us first construct our initial $F$ and $y$ (for $i = 0$). We define the MN $F$ with $m = 3$ as follows:

$$\forall z \in \mathbb{B}^n, \forall j \in [n], \begin{cases} f_j(z) = 0 & \implies \forall z' \in \Pi_{i=1}^n \{2 \cdot z_i, (2 \cdot z_i + 1)\}, F_j(z') = -1 \\ f_j(z) = 1 & \implies \forall z' \in \Pi_{i=1}^n \{2 \cdot z_i, (2 \cdot z_i + 1)\}, F_j(z') = 1 \end{cases}$$

We first show that $F$ is indeed a refinement of $f$. Let $z \in \mathbb{M}^n$ and $j \in [n]$ be arbitrary such that $F(z) = -1$ as the case of $F(z) = 1$ is symmetric. We want to show $\exists z' \in \beta(z)$ such that $f(z') = 0$.

Consider the state $z'$ defined as follows:

$$\forall j \in [n], z'_j = \frac{z_j - (z_j \bmod 3)}{3}$$

Surely such state is a binarization of $z$, $z' \in \beta(z)$. Moreover, $f(z') = 0$ as by definition of $F$, $f(z') \implies F(z) = 1$ leads to a contradiction.

Let us define $y = 3.x$: it is trivially a trace refinement of $x$ with the trace refinement function $\kappa : 0 \mapsto 0$.

We now iterate over $i \in \{1, \ldots, k\}$, adjusting $F$, $y, y', \ldots, y^{(l_i)}$ and $\kappa$ to ensure $y, y', \ldots, y^{(l_i)}$ is a trace refinement of $x, x', \ldots, x^{(i)}$. Moreover, we maintain that no transition increases any component value beyond 2 or decreases below 1 along $y', \ldots, y^{(l_i)}$ and ensure that $\forall j \in [n]$, $x_j^{(i)} = \searrow \implies y_j^{(\kappa(i))} = 1$ and $x_j^{(i)} = \nearrow \implies y_j^{(\kappa(i))} = 2$.

Let $\{e\} = \Delta(x^{(i-1)}, x^{(i)})$ and let $l_{i-1}$ denote the current length of the sequence of configurations $y, y', \ldots, y^{(l_{i-1})}$. We modify $F$ and extend $y, y', \ldots, y^{(l_{i-1})}$ based on the value of $x_e^{(i)}$:

- $x_e^{(i)} \notin \mathbb{B}$. Let us assume $x_e^{(i)} = \searrow$ without loss of generality, as the construction is symmetric for $x_e^{(i)} = \nearrow$.

  First, we extend $y, y', \ldots, y^{(l_{i-1})}$ based on $y_e^{(l_{i-1})}$:

  - $y_e^{(l_{i-1})} = 3$, $y^{(l_{i-1}+1)} := z \wedge y^{(l_{i-1}+2)} := z'$;
  - $y_e^{(l_{i-1})} = 2$, $y^{(l_{i-1}+1)} := z'$;

  where $z$ and $z'$ are equal to $y^{(l_{i-1})}$ but $z_e = 2$ and $z'_e = 1$. The trace refinement function is adjusted accordingly, $y_e^{(l_{i-1})} = 3 \implies \kappa : i \mapsto l_{i-1} + 2 = l_i$ and $y_e^{(l_{i-1})} = 2 \implies \kappa : i \mapsto l_{i-1} + 1 = l_i$.

  If $\forall 0 < j \leq l_i$, $y^{(j-1)} \xrightarrow[\text{a}]{F} y^{(j)}$, we are done. Otherwise, we modify $F_e(y^{(l_{i-1})}) := -1$ and, if necessary, also $F_e(z) := -1$. Since for any $j \in [n]$, $y_j^{(l_{i-1})} \in \{1, 2\}$ exactly when $x_j^{i-1} \notin \mathbb{B}$, the new $F$ is a refinement of $f$. $\forall 0 < j \leq l_i$, $y^{(j-1)} \xrightarrow[\text{a}]{F} y^{(j)}$ holds in the new $F$ as the $e$-th component never increases value beyond 2 along $y, y', \ldots, y^{(l_i)}$.

  Finally, $y, y', \ldots, y^{(l_i)}$ is indeed a trace refinement of $x, x', \ldots, x^{(i)}$ with $\kappa$:

  1. $\kappa$ being non-decreasing is guaranteed as $l_i > l_{i-1}$.
  2. $\kappa(0) = 0$ remains unchanged from the initial step and $\kappa(i) = l_i$ by definition.
  3. $0 < y_e^{(l_i)} = 1 < 3$ and the rest follows from the induction hypothesis.
  4. $F_e(y^{(\kappa(i-1))}) = F_e(y^{(l_{i-1})}) = -1$.

- $\pi_i(k) \in \mathbb{B}$. No change is made safe for the completion of the trace refinement function $\kappa : i \mapsto \kappa(i-1)$. $y, y', \ldots, y^{(l_{i-1})}$ being a trace refinement of $x, x', \ldots, x^{(i)}$ is trivial as the fourth point of Definition 2.7 is not applicable.

□

**C. Computational complexity.** We address the computational complexity of basic dynamical properties, as done in Sect. 1.D, but with the most permissive semantics.

First, remark that fixed points of the most permissive semantics are exactly the fixed points of the (a)synchronous semantics (for any configuration $x \in \mathbb{P}^n$, $\rho^f_{\text{mp}}(x) = \{x\} \Leftrightarrow x \in \mathbb{B}^n \wedge f(x) = x$). Therefore the complexity of deciding if a configuration $x$ is a fixed point is the same (NP-complete, Proposition 1).

***C.1. Theorem 3 (Reachability).*** Lemma 1 establishes that if there exists a sequence of most-permissive transitions from a configuration $x$ to a configuration $y$, then there exists a sequence of linear length linking the two configurations. Lemma 2 then states that searching for such a sequence requires exploring at most a quadratic number of transitions, which leads to Theorem 3 establishing the computational complexity for deciding reachability as in P for locally-monotonic BNs and in $\mathrm{P}^{\mathrm{NP}}$ (also known as $\Delta_2^P$) otherwise.

**Lemma 1.** *Given a BN $f$ of dimension $n$ and any configurations $x, y \in \mathbb{B}^n$, if $x \xrightarrow[\text{mp}]{f}{}^* y$, then there exists a sequence of at most $3n$ transitions $\xrightarrow[\text{mp}]{f}$ from $x$ to $y$. This sequence starts with at most $n$ and at least $|\Delta(x,y)|$ transitions of the form $\mathbb{B} \to \{\nearrow, \searrow\}$, then at most $n$ transitions of the form $\{\nearrow, \searrow\} \to \{\searrow, \nearrow\}$, and then at most $n$ transitions of the form $\{\nearrow, \searrow\} \to \mathbb{B}$.*

*Proof.* Let us consider any sequence of transitions $x \xrightarrow[\text{mp}]{f} w^1 \xrightarrow[\text{mp}]{f} \cdots w^k \xrightarrow[\text{mp}]{f} y$. Let us define the set of components which went through the state $\nearrow$ or $\searrow$ during this sequence of transitions, $\hat{I} := \{i \in [n] \mid \exists j \in [k], w^j \notin \mathbb{B}\}$.

Let us prove that there exists $\hat{z} \in \mathbb{P}^n$ with $\Delta(x, \hat{z}) = \hat{I}$ and $\forall i \in \hat{I}$, $\hat{z}_i \notin \mathbb{B}$, such that $x \xrightarrow[\text{mp}]{f}{}^* \hat{z}$ in $|\hat{I}|$ transitions. For each component $i \in \hat{I}$, we write $\nu(i)$ the smallest index $j \in [k]$ such that $w_i^j \neq \mathbb{B}$. Necessarily, for each $i \in \hat{I}$, $\exists z \in \gamma(w^{\nu(i)-1}) : f_i(z) \neq x_i$, identifying $w^0$ with $x$. The components in $\hat{I}$ can then be ordered as $\{i^1, \dots, i^{|\hat{I}|}\} = \hat{I}$ with $\nu(i^1) < \cdots < \nu(i^{|\hat{I}|})$. First, remark that $\nu(i^1) = 1$, hence $x \xrightarrow[\text{mp}]{f} z^1$ with $\Delta(x, z^1) = \Delta(w^{\nu(i^1)-1}, w^{\nu(i^1)}) = \{i^1\}$ and $z^1_{i^1} = w^{\nu(i^1)}_{i^1}$. Then, remark that $\gamma(w^{\nu(i^2)}) \subseteq \gamma(z^1)$, hence, $z^1 \xrightarrow[\text{mp}]{f} z^2$ with $\Delta(z^1, z^2) = \Delta(w^{\nu(i^2)-1}, w^{\nu(i^2)}) = \{i^2\}$ and $z^2_{i^2} = w^{\nu(i^2)}_{i^2}$. By induction, we obtain $x \xrightarrow[\text{mp}]{f}{}^* \hat{z}$. Remark that $\forall i \in \hat{I}$, $\hat{z}_i = \nearrow$ whenever $x_i = 0$ and $\hat{z}_i = \searrow$ whenever $x_i = 1$.

Now, let us consider the subset of components in $\hat{I}$ which are equal in $x$ and $y$, $\bar{I} := \{i \in \hat{I} \mid x_i = y_i\}$: for each of these components $i \in \bar{I}$, there exists $j' \in \{\nu(i), \dots, k\}$ such that $w_i^{j'} = \searrow$ whenever $x_i = y_i = 0$ and $w_i^{j'} = \nearrow$ whenever $x_i = y_i = 1$. By definition of $\hat{I}$ and $\hat{z}$, we obtain that $\gamma(w^{j'}) \subseteq \gamma(\hat{z})$. Therefore, there exists $\check{z} \in \mathbb{P}^n$ with $\Delta(\hat{z}, \check{z}) = \bar{I}$ and $\hat{z} \xrightarrow[\text{mp}]{f}{}^* \check{z}$ using $|\bar{I}|$ transitions. Finally, remark that $\check{z} \xrightarrow[\text{mp}]{f}{}^* y$ using $|\hat{I}|$ transitions.

In summary, $x \xrightarrow[\text{mp}]{f}{}^* \hat{z} \xrightarrow[\text{mp}]{f}{}^* \check{z} \xrightarrow[\text{mp}]{f}{}^* y$ in $|\hat{I}| + |\bar{I}| + |\hat{I}| \leq 3n$ iterations. □

**Lemma 2.** *Given a BN $f$ of dimension $n$ and any configurations $x, y \in \mathbb{B}^n$, deciding if $x \xrightarrow[\text{mp}]{f}{}^* y$ requires computing at most $\frac{n(n-1)}{2}$ transitions of $\xrightarrow[\text{mp}]{f}$; whenever $y$ belongs to an attractor, it requires as most $n$ transitions.*

*Proof.* Let us consider the following procedure with $L \subseteq [n]$, initially with $L = \emptyset$:

1. From $x$, apply only transitions of the form $\mathbb{B} \to \{\nearrow, \searrow\}$ to components $i \in [n] \setminus L$. Let us denote by $\hat{z}^L \in \mathbb{P}^n$ the (unique) reached configuration.
2. If $y \notin \gamma(\hat{z}^L)$, then $y$ is not reachable from $x$.
3. Otherwise, let us consider the components that cannot reach their value in $y$ from $\hat{z}^L$, $\bar{I}^L := \{i \in [n] \mid \hat{z}^L_i \notin \mathbb{B} \wedge \nexists z \in \gamma(\hat{z}^L), f_i(z) = y_i\}$:
   (a) If $\bar{I}^L = \emptyset$, then $\hat{z}^L \xrightarrow[\text{mp}]{f}{}^* y$.
   (b) Otherwise, repeat the procedure with $L := L \cup \bar{I}^L$.

Remark that this procedure can be iterated at most $n$ times, each of them computing at most $n - |L|$ transitions. Its correctness can be demonstrated as follows.

By Lemma 1, $x \xrightarrow[\text{mp}]{f}{}^* y$ if and only if there exists $L \subseteq [n]$ such that $y \in \gamma(\hat{z}^L)$ and $\bar{I}^L = \emptyset$. Notice that there is a unique $\subseteq$-minimal $L^*$ verifying $y \in \gamma(\hat{z}^{L^*})$ and $\bar{I}^{L^*} = \emptyset$: if $L^1$ and $L^2$ verify these properties, then so does $L^1 \cap L^2$.

Let us denote by $L^0, \dots, L^m$ the successive values of $L$ at the beginning of each iteration of the procedure ($L^0 = \emptyset$). We prove that $L^* = L^m$. Let us admit that $L^k \subseteq L^*$ with $k < m$. By construction, $\gamma(\hat{z}^{L^*}) \subseteq \gamma(\hat{z}^{L^k})$. Let us assume there exists $i \in \bar{I}^{L^k}$ and $i \notin L^*$. Then, $\hat{z}^{L^*} = \hat{z}^{L^k} \notin \mathbb{B}$, and there exists $z \in \gamma(\hat{z}^{L^*})$ with $f_i(z) = y_i$, which is a contradiction.

Whenever $y$ belongs to an attractor, $\bar{I}^\emptyset = \emptyset$. Indeed, remark that $\rho^f_{\text{mp}}(y) \subseteq \gamma(\hat{z}^\emptyset)$. Thus, if there exists a component $i \in \bar{I}^\emptyset$, then from any configuration $y' \in \rho^f_{\text{mp}}(y)$, $y \notin \rho^f_{\text{mp}}(y')$, which is a contradiction. Therefore, the procedure is executed only once, which involves computing at most $n$ transitions. □

Steps 1 and 3 of the procedure check for the existence of transitions in a most-permissive configuration, i.e., for the existence of a binary configuration compatible with it and such that the local function has a given value. This is exactly the SAT problem, which is NP-complete in the general case, and P whenever $f$ is locally-monotonic.

**Theorem 3.** *Given a BN $f$ of dimension $n$ and two configurations $x, y \in \mathbb{B}^n$, deciding if $y \in \rho^f_{\text{mp}}(x)$ is in P if $f$ is locally-monotonic, and in $P^{NP}$ otherwise.*

***C.2. Theorem 4 (Attractors).*** Attractors of the BN $f$ with the most permissive semantics match exactly with the *minimal trap spaces* of $f$ (4) (Proposition 8). Thus, Deciding if a given configuration $x$ belongs to an boils down to deciding if the smallest hypercube closed by $f$ and containing $x$ is minimal.

The fact that an attractor is necessarily an hypercube comes from the property that if two configurations lying on a diagonal of an hypercube are within the same attractor, then all adjacent configurations are within the attractor as well. This is illustrated by the following drawing, where boxed configurations belongs to a same attractor:

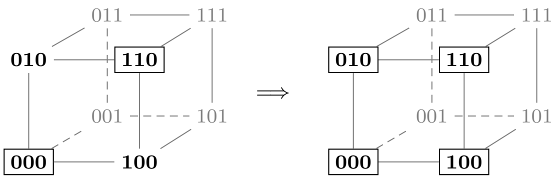

**Proposition 8.** *$A \subseteq \mathbb{B}^n$ is an attractor of $f$ with the most permissive semantics if and only if there exists a minimal hypercube $h \in (\mathbb{B} \cup \{*\})^n$ closed by $f$ such that $c(h) = A$.*

*Proof.* Let us consider a configuration $x \in A$, and let $h \in (\mathbb{B} \cup \{*\})^n$ be the smallest hypercube closed by $f$ containing $x$. Let us denote by $y \in c(h)$ the configuration of this hypercube which is the most distant from $x$: $\forall i \in [n]$, $y_i = \neg x_i$ whenever $h_i = *$, otherwise $y_i = x_i$. According to the previous section, remark that $y$ is reachable from $x$; thus by attractor hypothesis, $x$ is also reachable from $y$. Now remark that the smallest hypercube closed by $f$ and containing $y$ is the same $h$ (otherwise $h$ would not be closed). Thus, for any component $i \in [n]$ which is free in $h$ ($h_i = *$), there exists a configuration $z \in c(h)$ such that $f_i(z) = 0$ and a configuration $z' \in c(h)$ such that $f_i(z') = 1$. Therefore, any configuration of the hypercube $h$ is reachable from $x$. Finally, notice that if there exists a configuration $y \in c(h)$ reachable from $x$, but where $x$ does not belong to the smallest hypercube closed by $f$ and containing $y$, then $x$ is not reachable from $y$, and thus does not belong to any attractor. □

**Theorem 4.** *Given a BN $f$ of dimension $n$ and a configuration $x \in \mathbb{B}^n$, deciding if $x$ belongs to an attractor of $f$ with the most permissive semantics is in coNP whenever $f$ is locally monotonic, and in $coNP^{coNP}$ otherwise.*

*Proof.* Consider IS-NOT-CLOSED$(f, h)$ the problem of deciding if the given hypercube $h$ is *not* closed by $f$: it is equivalent to deciding if there exists component $i \in [n]$ with $h_i \neq *$ and $z \in c(h)$ such that $f_i(z) \neq h_i$, which is NP-complete in general, and P whenever $f$ is locally monotonic. Then, the complementary problem IS-CLOSED$(f, h)$ is in coNP in the general case and in P in the locally-monotonic case.

Consider IS-NOT-MINIMAL$(f, h)$ the problem of deciding if the hypercube $h$ closed by $f$ is *not* minimal: it can be solved by deciding wherever there exists an hypercube $h'$ which is strictly included in $h$ and which is closed by $f$, which is at most $\text{NP}^{\text{IS-CLOSED}}$. Thus, the complementary problem IS-MINIMAL$(f, h)$ is in $\text{coNP}^{\text{IS-CLOSED}}$, i.e., $\text{coNP}^{\text{coNP}} = \Pi^{\text{P}}_2$ in the general case and coNP in the locally-monotonic case. □

 

### 3. Case studies

**A. Code.** A simple implementation of reachability and attractor computations for Most Permissive Boolean Networks is available at https://github.com/pauleve/mpbn. It relies on Answer-Set Reprogramming (5) and the solver `clingo` (6) which offers features such as minimal model enumeration.

The computational analyzes have been performed within the CoLoMoTo environment (7) and can thus be reproduced using the provided notebook files within the Docker image `colomoto/colomoto-docker:2020-03-19`:

Using Python (https://python.org), execute the following command in a terminal:

```
sudo pip install -U colomoto-docker # you may have to use pip3
colomoto-docker -V 2020-03-19
```

Alternatively, you can run the image directly with Docker (https://docker.com):

```
docker run -it --rm -p 8888:8888 colomoto/colomoto-docker:2020-03-19
```

and then open your webbrowser to https://localhost:8888. See https://colomoto.org/notebook for detailed instructions.

The notebook files (with `.ipynb` extension) used in the next sections can be downloaded from http://doi.org/10.5281/zenodo.3719097 and then be uploaded and executed within the Jupyter web interface.

**B. Models of differentiation processes from literature.** We show on several case studies from literature that MPBNs, although potentially predicting more behaviors than asyncrhonous BNs, are still stringent enough to predict cell fate decision processes, i.e., absence of attractor reachable from specific configurations or with specific perturbations.

***B.1. Tumour invasion model by Cohen et al. 2015.*** (8) This BN (32 components, Fig. S2) models cellular decision processes involved in tumour invasion, with attractors related to apoptosis, cell cycle arrest, and various stages leading to metastasis. The study analyzed the reachability of these attractors from a set of initial conditions and subject to different mutations. We reproduced the analysis of reachable attractor from the same initial condition in the wild-type model and with the sol NICD gain of function and the combined NICD gain of function and p53 loss of functions. In each case, MPBNs repart the same reachable attractors than the original analysis with fully asynchronous BNs, the later double mutant leading to the loss of capability to reach apoptotic attractors.

The details of the analysis can be found in the notebook "*MPBN applied to Tumour invasion model by Cohen et al. 2015.ipynb*", which can be visualized online at doi:10.5281/zenodo.3936123.

***B.2. T-cell differentiation model by Abou-Jaoudé et al. 2015.*** (9) This multivalued network (101 components, Fig. S3) models reprogramming capabilities across different T-cell types. T-cell types are modeled by attractors where pre-determined markers have a fixed value. The authors also pre-determined input conditions of the signaling pathways, which should be associated to the inevitable reachability of specific T-cell types. The original study then analyzed which input conditions can stir (inevitably) the system from any state (not only in attractors) matching with the markers of a T-cell type to an attractor matching a different T-cell type. This results in a so-called reprogramming graph, where nodes are the T-cell subtypes, and directed edges show the possible trans-differentiation, parameterized by the input condition triggering it.

Because of the size of the model, the study had to rely on approximations through model reduction, and use symbolic model-checkers to avoid computing explicitly the attractors of the model.

Employing MPBNs after a booleanization (10) of the full-size model, we are able to explicitly compute all the attractors with the MP semantics in each of the defined input conditions. It turns out that in most of them, all the attractors are fixed points, except in APC and proTh1 conditions which both have one complex attractor. Fixed points are invariant in MPBNs and (fully) asynchronous BNs, and to each complex MP attractor corresponds at least one complex asynchronous attractor.

The details of the analysis can be found in the notebook "*MPBN applied to T-Cell differentiation model by Abou-Jaoudé et al. 2015.ipynb*", which can be visualized online at doi:10.5281/zenodo.3936123.

**C. Scalability.** The theoretical complexity gain brought by the Most Permissive semantics has a drastic impact for the analysis of large BNs. We illustrate it both on large networks from literature, and on very large networks (up to 100,000 components) generated randomly. Computation times are obtained on an Intel(R) Xeon(R) E-2124 CPU @ 3.30GHz.

***C.1. Networks from literature.*** In the two analysis of the previous section, the computation of reachable attractors takes less than 10ms on the Tumour invasion model (32 components) and less than 100ms on the T-cell differentiation model (104 components).

We additionally performed reachable attractor computations on the **Bladder tumorigenesis model by Remy et al. 2015** (11), as it served as benchmark for evaluating simulations methods for devising reachable attractors and their propensity in (12). The network has 35 components. The computation of reachable attractors in diverse settings are performed in less than 10ms, whereas simulations of asynchronous BNs were reported taking from 10s to 700s, and possibly not successful. Note however that contrary to the simulation methods, we are not able to compute nor estimate the propensity of reachable attractors. Nevertheless, the enumeration is guaranteed to be complete. In this application, the number of attractors of Most Permissive semantics is the same as the reported with fully asynchronous semantics.

The details of the analysis can be found in the notebook "*MPBN applied to Bladder Tumorigenesis by Remy et al 2015.ipynb*" which can be visualized online at doi:10.5281/zenodo.3936123.

***C.2. Very large networks.*** We generated random influence graphs (13) with scale-free structure (14) with a number of components ranging from 1,000 to 100,000 with in-degrees up to 1,400. We then applied the inhibitor dominant rule to assign a Boolean function to each component: the activation occurs only in configurations whenever no inhibitor are active and at least one activator is active. The following computations are measured: computation of 1 attractor; enumeration of (at most) 1000 attractors; and enumeration of (at most) 1000 attractors reachable from random initial configurations.

The details of the analysis can be found in the notebook "Scalability on large random BNs.ipynb" which can be visualized online at doi:10.5281/zenodo.3936123.

Fig. S4 summarizes the results and shows that the computation of reachable attractors take a fraction of a second with 1,0000 components, less than 2 seconds with 10,000, and less than 50 seconds with 100,000 components. The computation times exclude the time for parsing the input text file (up to 20s for larger networks).

### 4. I3-FFL network motif

The incoherent feed-forward loop of type 3, I3-FFL (15, Fig. S1) describes a simple interaction motif where an input node 1 directly inhibits the output 3, but indirectly activates it via node 2. This network can then be excited by a signal on node 1. According to theoretical studies (16, 17) employing quantitative models and experimental data from synthetically designed circuits (18), the following behaviors are observed under some range of kinetics parameters:

- with no signal, the system stays steady showing no activity;
- switching on the signal enables observing a transient activation of the output (node 3) before reaching a stable state where nodes 1 and 2 are active and node 3 inactive.

We show in the main text that synchronous and asynchronous analysis of the corresponding BN fails to predict this behavior, whereas the MP interpretation enable recovering it.

One may wonder whether there exist different BN matching the network motif and the observed behavior. We employed the in-development software BoNesis (19) which provides an exhaustive identification of all BNs compatible with a given influence graph and dynamical properties under the MP semantics.

It results that the BN depicted in Fig. S1 is the only compatible BN under the MP semantics. Notice that the input properties refer to the existence of two fixed points and of specific trajectories. Therefore, because MPBNs have the same fixed points as (a)synchronous BNs, the set of MPBNs compatible with the dynamical properties includes all the BNs compatible with (a)synchronous semantics.

Therefore, an (a)synchronous Boolean analysis would conclude that the network motif has to be revised in order to reproduce the observed behavior, whereas it is actually sufficient.

The details of the analysis can be found in the notebook "I3FFL - compatible MPBNs.ipynb" which can be visualized online at doi:10.5281/zenodo.3936123.

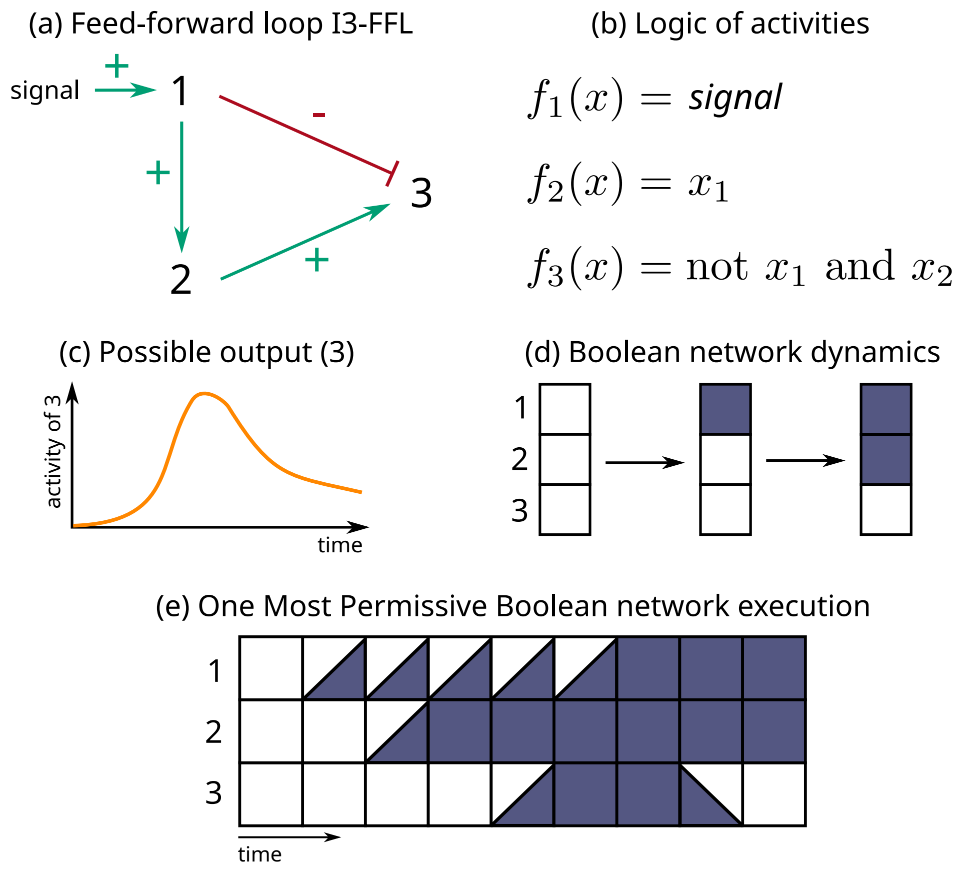

**Fig. S1.** Incoherent feed-forward loop of type 3 (a) and its associated Boolean logic for nodes activities (b). Whereas theoretical and experimental studies show a possible activation of the output (c), usual Boolean networks analysis cannot predict this transient behavior (d). Conversely, the Most Permissive interpretation can reproduce this behavior, (e) showing one of its possible execution. A configuration is represented by a pile of 3 squares, where the top square represents the state of the first component, and so forth. A white square represent the inactive (0) state; a blue square represents the active (1) state; the increasing state $\nearrow$ is represented by a top-right white and bottom-left blue square, and decreasing state $\searrow$ by a top-right blue and bottom-left white square.

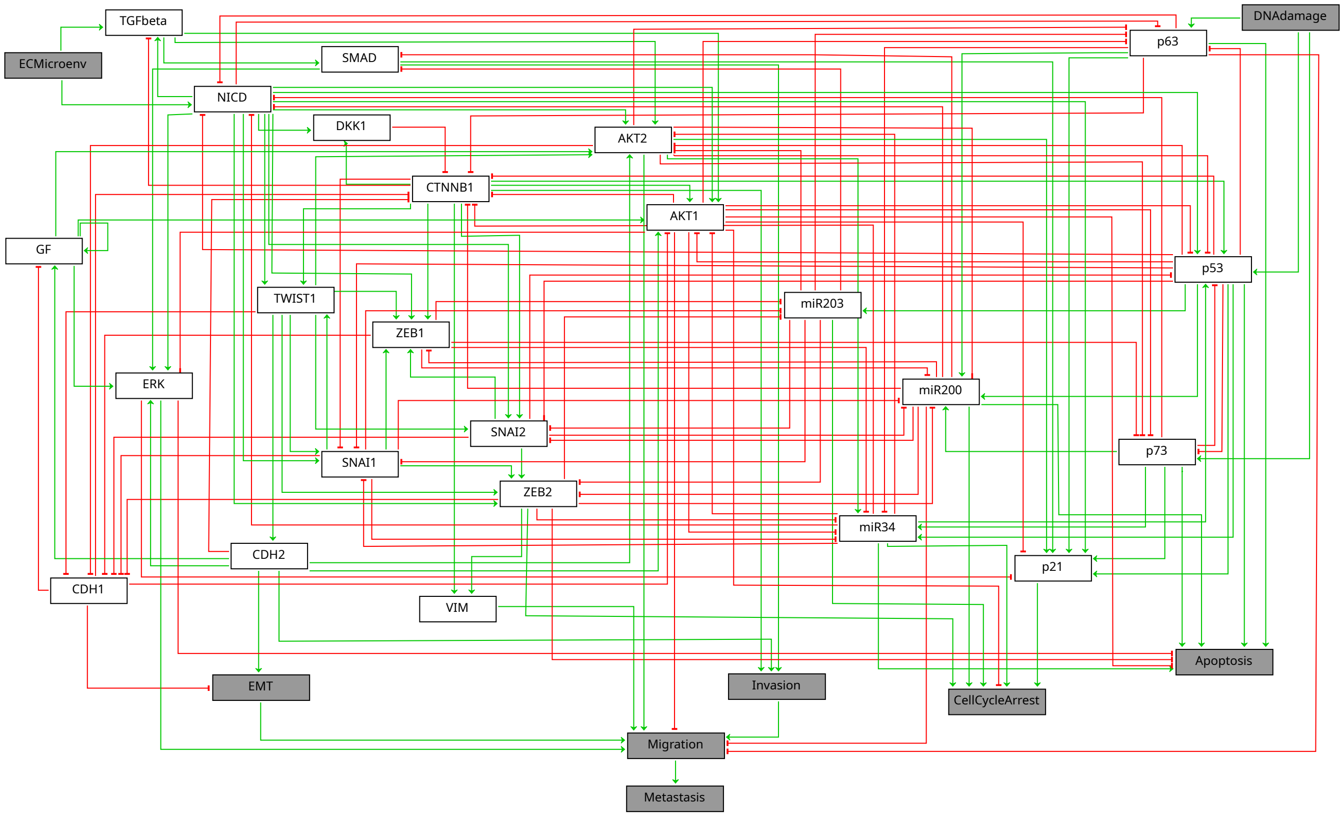

**Fig. S2.** Influence graph of the tumour invasion model by Cohen et al. (2015)

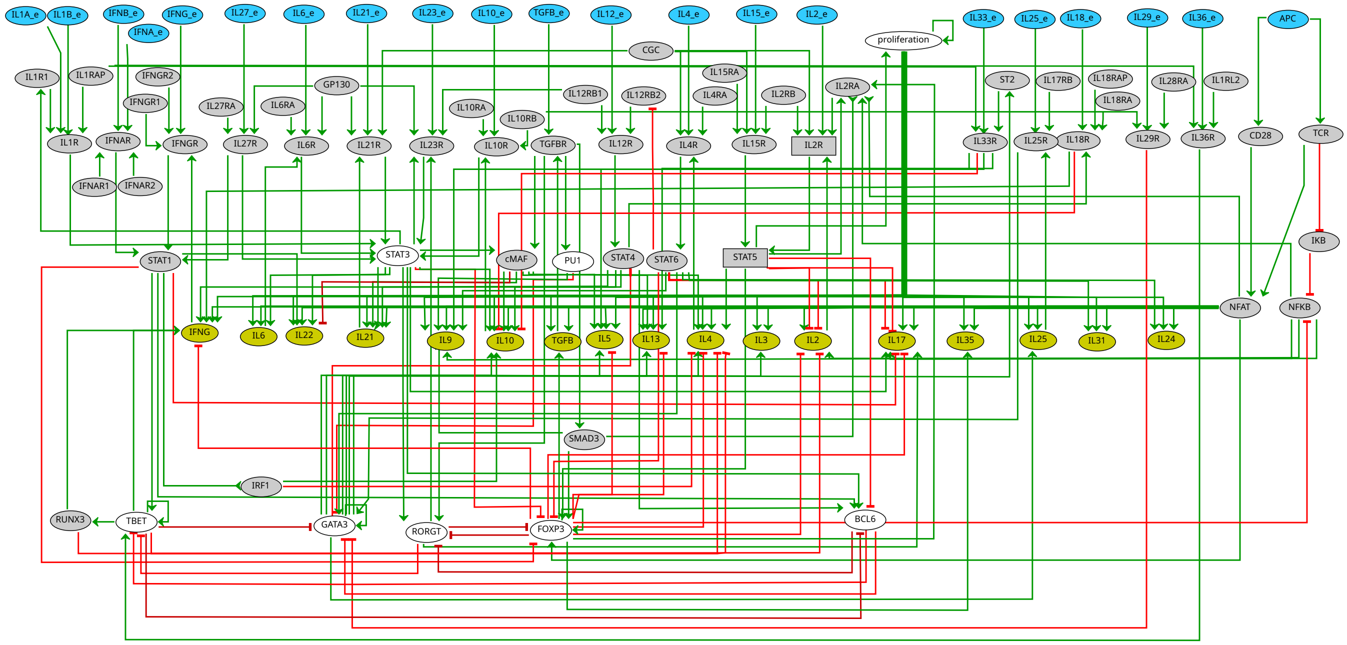

**Fig. S3.** Influence graph of the T-cell differentiation model by Abou-Jaoudé et al. (2015)

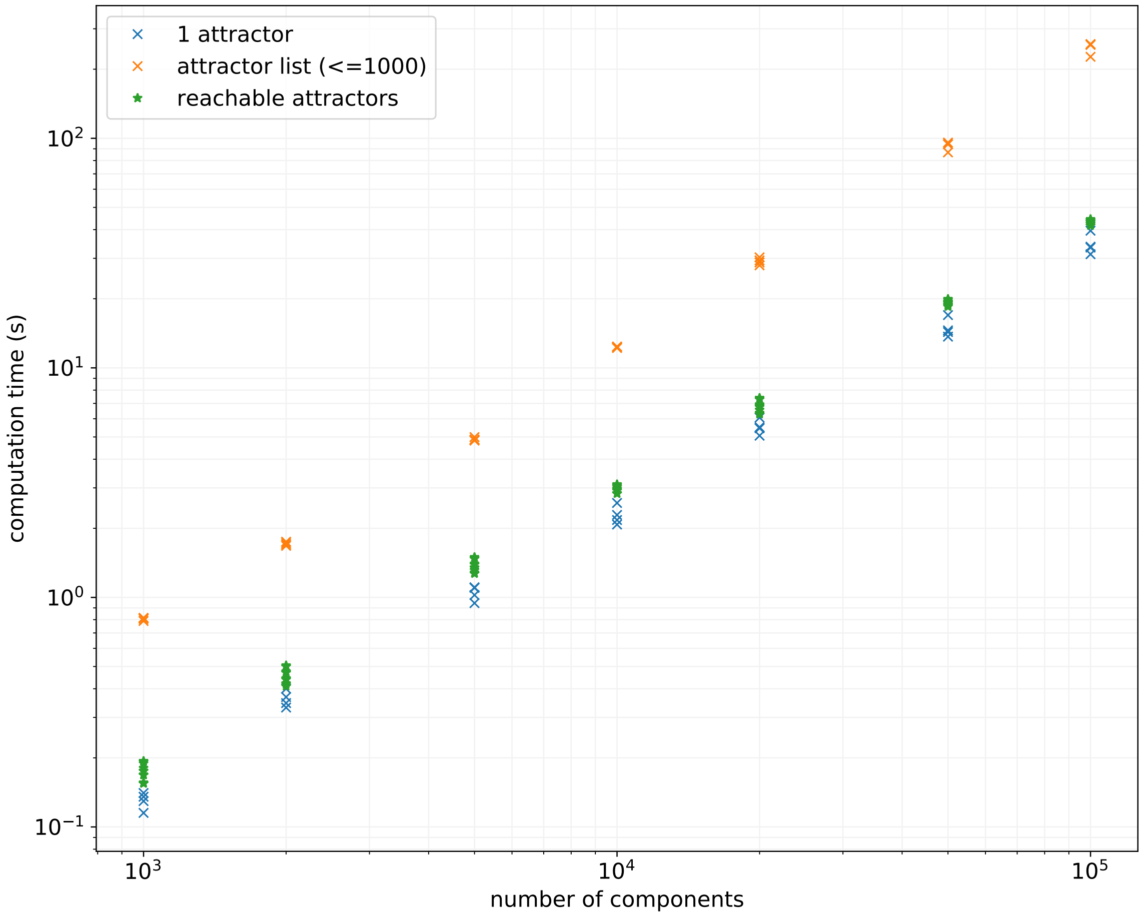

**Fig. S4.** Computation times on a 3GHz CPU obtained using `mpbn` software tool on BNs generated with random scale-free influence graph for the computation of a single attractor, the enumeration of 1,000 attractor, and the computation of attractors reachable from random initial conditions. See notebook “Scalability on large random BNs.ipynb” available at doi:10.5281/zenodo.3936123.
